# Id4 eliminates the pro-activation factor Ascl1 to maintain quiescence of adult hippocampal stem cells

**DOI:** 10.1101/426015

**Authors:** Isabelle Blomfield, Brenda Rocamonde, Maria del Mar Masdeu, Eskeatnaf Mulugeta, Stefania Vaga, Debbie L. C. van den Berg, Emmanuelle Huillard, François Guillemot, Noelia Urbán

## Abstract

Quiescence is essential for the long-term maintenance of adult stem cells and tissue homeostasis. However, how stem cells maintain quiescence is still poorly understood. Here we show that stem cells in the dentate gyrus of the adult hippocampus actively transcribe the pro-activation factor Ascl1 regardless of their activation state. We found that the inhibitor of DNA binding protein Id4 suppresses Ascl1 activity in neural stem cell cultures. Id4 sequesters Ascl1 heterodimerisation partner E47, promoting Ascl1 protein degradation and neural stem cell quiescence. Accordingly, elimination of Id4 from stem cells in the adult hippocampus results in abnormal accumulation of Ascl1 protein and premature stem cell activation. We also found that multiple signalling pathways converge on the regulation of Id4 to reduce the activity of hippocampal stem cells. Id4 therefore maintains quiescence of adult neural stem cells, in sharp contrast with its role of promoting the proliferation of embryonic neural progenitors.

## INTRODUCTION

Tissue stem cells must maintain their long-term activity while minimising the accumulation of genetic and metabolic damage. In several adult tissues, stem cells can remain inactive for long periods of time in a state of quiescence. Specific stimuli promote the exit from quiescence of different types of adult stem cells, such as hypoxia for stem cells of the carotid body or muscle injury for satellite stem cells (Dumont et al., 2015; Sobrino et al., 2018). Regulation of the transit between the quiescent and active compartments is essential to maintain an appropriate pool of stem cells able to sustain tissue homeostasis and provide an adequate response to insults over the lifespan of the organism. An excessive retention of stem cells in the quiescent compartment would not produce enough differentiated progeny to maintain functionality, as happens for instance during aging (Garcia-Prat et al., 2016; Leeman et al., 2018). On the other hand, excessive stem cell activity would eventually result in stem cell exhaustion, also leading to loss of functionality (Castilho et al., 2009; Ho et al., 2017). Quiescence is also an essential property of cancer stem cells that allows them to evade immune surveillance and results in resistance to treatment (Agudo et al., 2018). Despite their relevance for the fields of tissue repair, aging and cancer biology, the mechanisms regulating quiescence in adult stem cells are still largely unknown.

In the adult brain, neural stem cell (NSC) populations in the subependymal zone (SEZ) of the lateral ventricles and in the dentate gyrus (DG) of the hippocampus, generate new neurons and glia that integrate into pre-existing neuronal networks (Bond et al., 2015; Lim and Alvarez-Buylla, 2016). In both regions, a large fraction of stem cells is quiescent. Extensive work has led to the identification of extracellular signals present in the SEZ and DG niches that regulate quiescent and active states (Choe et al., 2015; Silva-Vargas et al., 2013). Notch, BMP4 and the neurotransmitter GABA have been shown to maintain stem cell quiescence while Wnt and Shh are thought to promote stem cell activity (Bao et al., 2017; Choe et al., 2015; Engler et al., 2018; Imayoshi et al., 2010; Lie et al., 2005; Mira et al., 2010; Petrova et al., 2013; Qu et al., 2010; Song et al., 2012). However, little is known of the cell intrinsic machinery that NSCs employ to adjust their activity to the different signals received from the niche. Ascl1 is one of the few intrinsic regulators of neural stem cell quiescence described so far. Ascl1 is a basic-helix-loop-helix (bHLH) transcription factor that is present in the adult hippocampus in a fraction of dividing stem cells and intermediate progenitors. Loss of Ascl1 completely blocks the activation of adult hippocampal stem cells, inhibits the generation of new neurons and prevents the depletion of the stem cell pool over time (Andersen et al., 2014). Ascl1 may therefore determine the balance between quiescence and activity of hippocampal NSCs. Indeed, stabilisation of Ascl1 protein by inactivation of the E3 ubiquitin ligase Huwe1 resulted in over-proliferation of hippocampal stem cells and prevented their return to quiescence (Urban et al., 2016). However, Huwe1 inactivation was not sufficient to trigger the activation of quiescent stem cells, indicating that additional mechanisms maintain the quiescent state of hippocampal stem cells.

The Id (inhibitor of differentiation/DNA binding) proteins are known inhibitors of bHLH transcription factors such as Ascl1 (Ling et al., 2014) (Imayoshi and Kageyama, 2014). Id proteins contain a conserved HLH domain with which they dimerise with bHLH proteins. However they lack a DNA binding domain and therefore prevent bHLHs with which they interact from binding DNA and other bHLH factors (Benezra et al., 1990). In mammals, the Id family comprises four genes: *Id1*-*4*. *Id1*-*3* are co-expressed in multiple tissues during development, and have been shown to promote stemness and proliferation in different systems, including in hematopoietic stem cells and in stem cells of the adult SVZ (Niola et al., 2012; Singh et al., 2018). In contrast, *Id4* expression rarely overlaps with that of the other *Ids* during development and it is generally thought to have an opposing function to Id1-3 (Patel et al., 2015). For instance in the context of cancer, Id4 has been shown to behave as a tumour suppressor, while other Id proteins are considered oncogenes (Patel et al., 2015; Ruzinova and Benezra, 2003; Sikder et al., 2003).

Here we show that Ascl1 mRNA is expressed by hippocampal stem cells independently of their proliferative state, but that only active stem cells reach significant levels of Ascl1 protein. This non-transcriptional regulation of Ascl1 is recapitulated in hippocampal stem cell cultures in vitro, where the quiescence-inducing factor BMP4 has no effect on Ascl1 mRNA expression but is sufficient to lower down Ascl1 protein levels. We performed a gene expression screen in these cells and found that Id4 is strongly induced in quiescent NSCs. Id4 sequesters the Ascl1 heterodimerisation partner E47 and the resulting Ascl1 monomers, which are unable to bind DNA, are rapidly degraded by the proteasome. Therefore, Id4 blocks the pro-activation transcriptional program driven by Ascl1 and keeps stem cells quiescent. Indeed, elimination of Id4 from the adult brain resulted in increased Ascl1 protein levels in stem cells of the hippocampus and in their rapid entry into the cell cycle. The role of Id4 in the regulation of hippocampal stem cells is unique among Id genes since gain- or loss-of-function of Id1, the only other family member expressed in a significant number of hippocampal stem cells, did not affect Ascl1 protein levels or hippocampal stem cell proliferation. Id4 is induced by BMP4 in NSC cultures and has been shown to be regulated by both BMP and Notch signals in embryonic neural progenitors. However, Id4 expression was maintained in vivo in the absence of both BMP and Notch signalling, indicating that multiple niche signals converge on the induction of Id4 to provide robustness to the quiescent state.

## RESULTS

### Ascl1 is transcribed in quiescent stem cells of the hippocampus

To investigate how Ascl1 expression is regulated in hippocampal NSCs, we first assessed the transcriptional activity of the *Ascl1* locus using the *Ascl1^KIGFP^* mouse reporter line, in which the GFP reporter replaces the *Ascl1* coding sequence and marks cells that transcribe the *Ascl1* gene (Leung et al., 2007). Hippocampal stem cells in vivo will be called hereafter radial glia-like cells (RGLs) while hippocampal stem cells in culture will be called NSCs. We identified RGLs by their expression of glial fibrillary acidic protein (GFAP), localisation of their nucleus in the subgranular zone of the DG and presence of a radial process extending towards the molecular layer. We found that 82.3±3.8% of all RGLs were positive for GFP in the hippocampus of P70 *Ascl1^KIGFP^* mice and therefore transcribed *Ascl1*. In contrast, only 1.9±0.3% of these cells expressed Ascl1 protein at a level detected with anti-Ascl1 antibodies (Fig. 1A, 1B). Notably, 83.8±4.1% of the RGLs that did not express Ki67 and were therefore quiescent expressed GFP, indicating a transcriptionally active *Ascl1* locus (Fig. 1C and 1D). Moreover, GFP immunolabeling intensity was comparable in active Ki67+ RGLs and quiescent Ki67-RGLs (Fig. 1E). We confirmed the presence of *Ascl1* transcripts at similar levels in quiescent and active RGLs using single molecule in situ hybridization (Fig. 1F, G). These results show that, unexpectedly, *Ascl1* is already expressed in quiescent RGLs and that NSC activation is not accompanied by the induction or marked upregulation of *Ascl1* transcription. The finding that quiescent and proliferating RGLs transcribe the *Ascl1* gene at comparable levels but only proliferating RGLs express detectable levels of Ascl1 protein, indicates that Ascl1 protein expression in quiescent hippocampal RGLs is regulated by a non-transcriptional mechanism.

**Figure 1.**
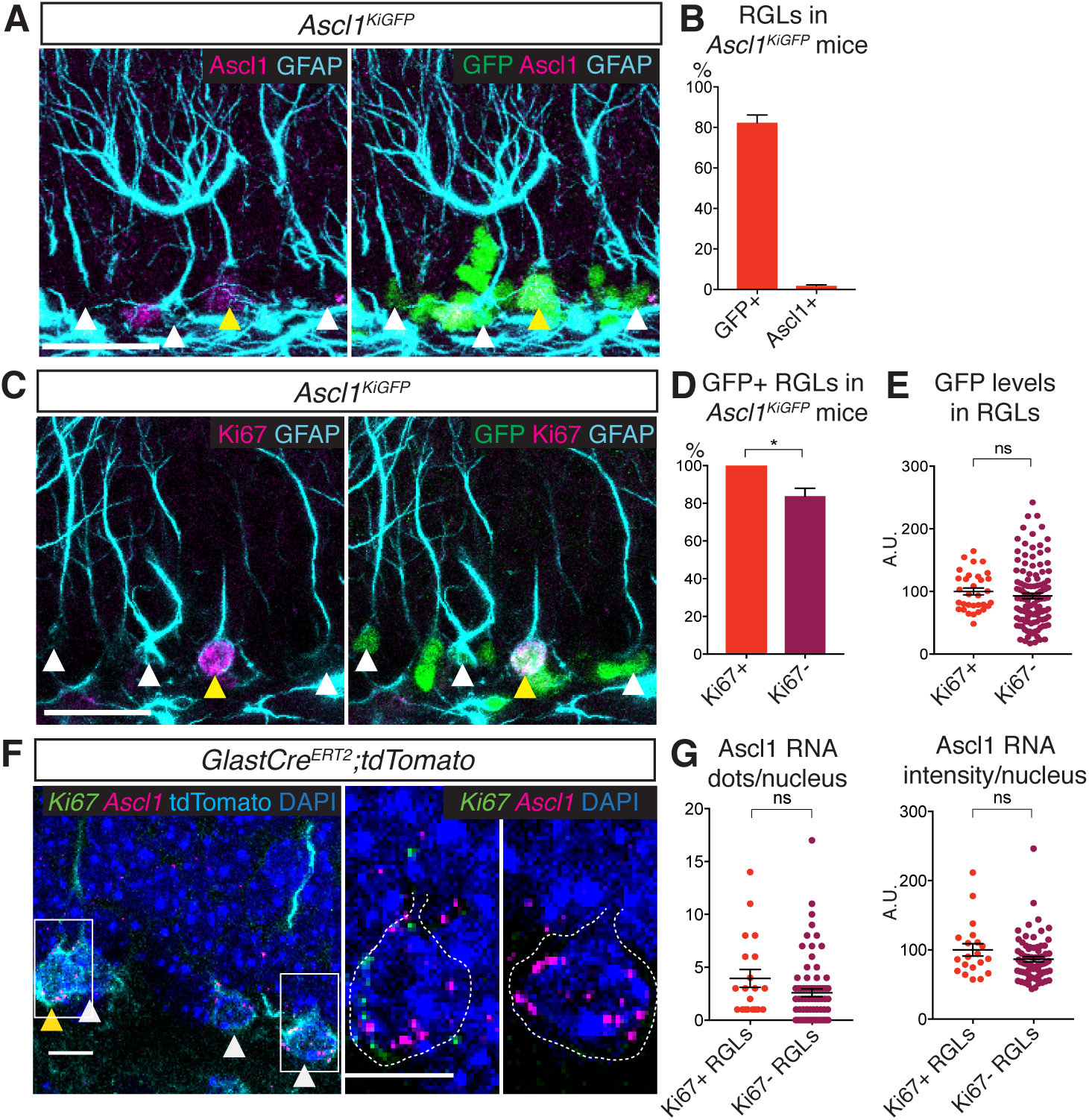
Ascl1 is transcribed in both quiescent and proliferating hippocampal stem cells. (A) Immunolabeling for GFP, Ascl1 and GFAP in the subgranular zone (SGZ) of the dentate gyrus (DG) of *Ascl1^KIGFP^* reporter mice. White arrows indicate GFP+Ascl1-RGLs; yellow arrows indicate GFP+Ascl1+ RGLs. Scale bar, 30μm. (B) Quantification of the data shown in (A). The widespread GFP expression indicates that Ascl1 is transcribed in most RGLs (radial GFAP+ cells) in *Ascl1^KIGFP^* mice while Ascl1 protein is only detectable in a small fraction of RGLs. n=3. (C) Immunolabeling for GFP, Ki67 and GFAP in the SGZ of the DG of *Ascl1^KIGFP^* reporter mice. White arrows indicate GFP+Ki67-RGLs; yellow arrows indicate GFP+Ki67+ RGLs. Scale bar, 30μm. (D, E) Quantification of the data in (C). Most quiescent (Ki67-) RGLs express GFP and therefore transcribe *Ascl1* (D) and the levels of GFP are not significantly different in quiescent and proliferating RGLs (E), indicating that Ascl1 is transcribed uniformly in the two RGL populations. p values, p>0.05 (ns). n=3. (F) RNA *in situ* hybridization by RNAscope^®^ with an Ascl1 probe (magenta) and a Ki67 probe (green) and immunolabeling for tdTomato to mark RGLs in the SGZ of the DG. To label RGLs with tdTomato, Glast-CreERT2;tdTomato mice were injected once at P60 with 4-hydroxytamoxifen, and analysed 48h later. White arrows indicate RGLs positive for Ascl1 RNA staining; yellow arrows show RGLs positive for both Ascl1 and Ki67 RNA. Magnifications of the RGLs marked by white boxes are shown on the right, highlighting an RGL positive for both Ascl1 and Ki67 RNA, and an RGL positive for only Ascl1 RNA. Dotted lines show the outline of the tdTomato signal. Scale bar, 10μm. n=5. (G) Quantification of the data in (F). Ascl1 transcripts are found at a similar level in quiescent (Ki67-) and proliferating (Ki67+) RGLs. Note the high variability in the levels of Ascl1 mRNA, which could be a reflection of the oscillatory nature of Ascl1 expression (Imayoshi et al., 2013). p values, p>0.05 (ns). n=5. Data are presented as mean +/- SEM, in this figure and all subsequent figures.

### Ascl1 is regulated post-translationally in quiescent NSC cultures

To investigate the mechanism regulating Ascl1 protein levels in quiescent hippocampal stem cells, we used an established cell culture model of NSC quiescence (Martynoga et al., 2013; Mira et al., 2010; Sun et al., 2011). The signalling molecule BMP4 has been shown to contribute to the maintenance of NSC quiescence in the hippocampus (Bonaguidi et al., 2008; Mira et al., 2010). BMP4 is also able to induce a reversible state of cell cycle arrest in embryonic stem cell-derived NSC cultures (Martynoga et al., 2013). Similarly, we found that NSCs originating from the adult hippocampus and maintained in culture in the presence of FGF2 stopped proliferating and entered quiescence when exposed to BMP4 (Fig. S1A-D). RNA sequencing analysis revealed that 1839 genes were differentially expressed between NSCs in proliferating and quiescent states (Fig. S1E, F). Ascl1 RNA levels were not significantly different between these two conditions as verified by QPCR (Fig. 2A). The intensity of GFP in cultured hippocampal NSCs derived from *Ascl1^KIGFP^* mice was also comparable in proliferating and quiescent conditions (2B, C), as for GFP expression in the hippocampus of *Ascl1^KIGFP^* mice (Fig. 1E). In contrast, Ascl1 protein levels were strongly reduced in BMP-treated quiescent NSCs (Fig. 2D-F), resembling the absence of Ascl1 protein in quiescent hippocampal RGLs in vivo (Fig. 1B). Treatment of quiescent NSCs with proteasome inhibitors significantly increased the levels of Ascl1 protein, suggesting that Ascl1 mRNA is translated in quiescent NSCs but Ascl1 protein is rapidly degraded in a proteasome-dependent manner (Fig. 2G). We previously showed that Ascl1 protein is targeted for proteasomal degradation by the E3 ubiquitin ligase Huwe1 in proliferating hippocampal NSC cultures (Urban et al., 2016). We therefore asked whether Huwe1 is also responsible for the degradation of Ascl1 in quiescent NSCs. We found that Ascl1 protein was similarly reduced in *Huwe1* mutant and control NSC cultures upon addition of BMP4 (Fig. S1G, H), indicating that a *Huwe1*-independent mechanism prompts the down-regulation of Ascl1 protein in quiescent cultured NSCs. This is in agreement with our prior observation that *Huwe1* is required for the return to quiescence of proliferating RGLs but not for the maintenance of RGL quiescence in mice (Urban et al., 2016). Together, these results establish BMP-treated NSC cultures as an appropriate model to characterise the mechanisms controlling Ascl1 protein levels in quiescent hippocampal stem cells.

**Figure 2.**
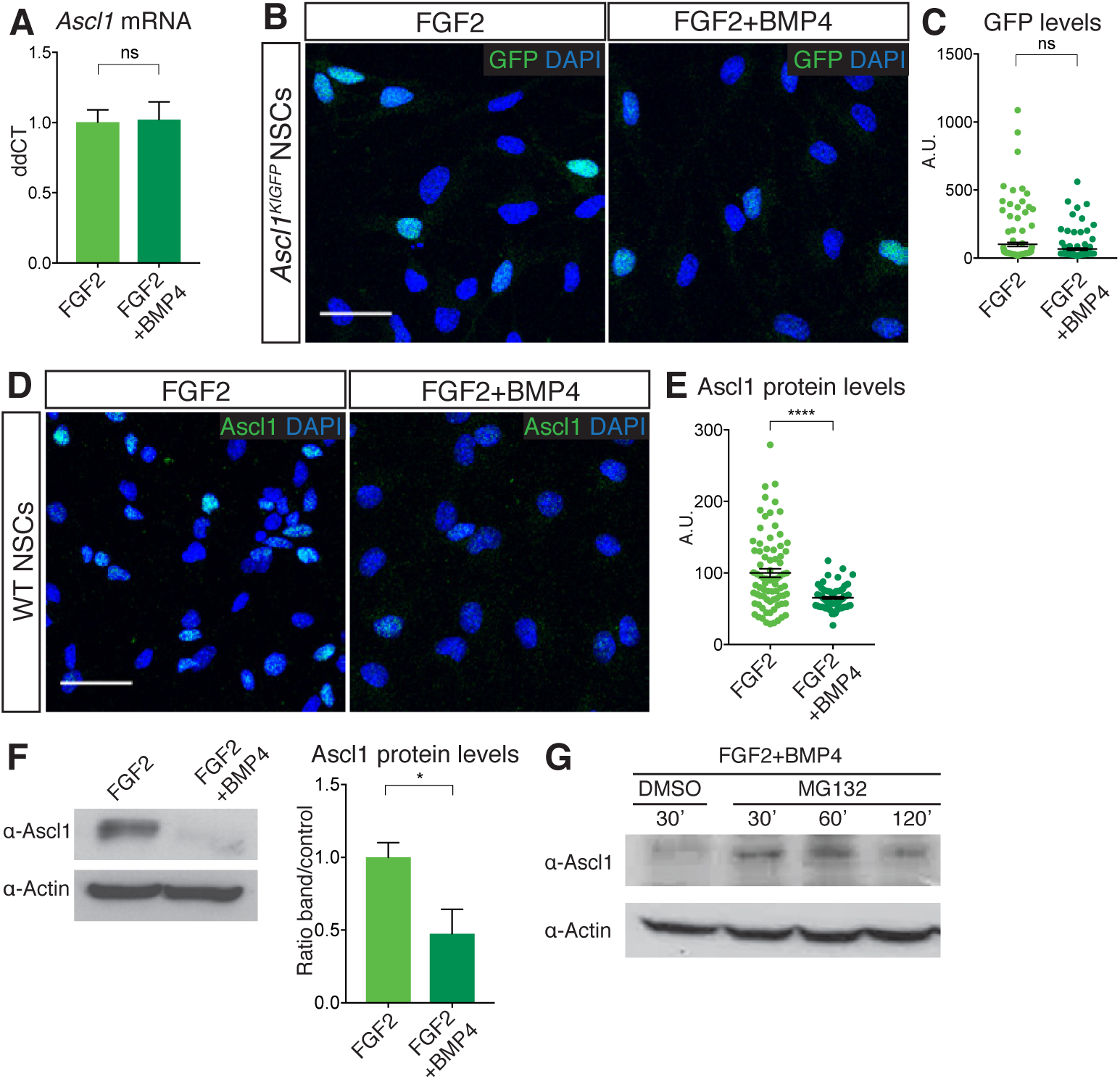
Ascl1 is regulated post-translationally in quiescent NSC cultures. (A) Transcript levels for *Ascl1* in hippocampus-derived NSC cultures treated with FGF2 alone (proliferating NSCs) or FGF2 and BMP4 (quiescent NSCs) analysed by QPCR. Ascl1 mRNA levels are unchanged in FGF2+BMP4-treated, quiescent NSCs. p>0.05 (ns), n=3. (B) Immunolabeling for GFP and DAPI staining in FGF2 and FGF2+BMP4-treated NSC cultures originating from *Ascl1^KIGFP^* mice. Scale bar, 30μm. (C) Quantification of the data in (B). GFP, which reports transcription of the Ascl1 gene, is expressed at similar levels in proliferating and quiescent NSCs. p>0.05 (ns). The data show one representative experiment, n=3. (D) Immunolabeling for Ascl1 and DAPI staining in FGF2- and FGF2+BMP4-treated NSC cultures. Scale bar, 30μm. (E) Quantification of the data in (D). Ascl1 levels are high in many proliferating NSCs and not detectable in most quiescent NSCs. The heterogeneity of Ascl1 expression in proliferating NSCs most likely reflects its oscillatory behaviour. p<0.0001 (****). n=3. (F) Western blot analysis and quantification of Ascl1 in FGF2- and FGF2+BMP4-treated NSCs. BMP4 suppresses Ascl1 protein expression. p<0.05 (*). n=3 (G) Western blot analysis of Ascl1 in FGF2+BMP4-treated NSCs after treatment with the proteasome inhibitor MG132 for different durations or with DMSO vehicle as a control. Ascl1 can be detected after proteasome inhibition in quiescent NSCs. See also Figure S1.

### Id4 is highly expressed in quiescent hippocampal stem cells in culture and in vivo

We screened our RNA-Seq data for potential Ascl1 inhibitory factors induced in quiescent conditions (Fig. S2). The four *Id genes Id1*-*4* were strongly induced in quiescent NSC cultures (Fig. S2A) and *Id3* and *Id4* are also enriched in quiescent hippocampal RGLs in vivo (Shin et al., 2015). Moreover, Id proteins have previously been shown to sequester E proteins, the dimerisation partners of Ascl1, resulting in the rapid degradation of Ascl1 monomers (Shou et al., 1999; Vinals et al., 2004). Id factors are therefore strong candidates to regulate Ascl1 post-translationally in hippocampal stem cells. Although the transcripts for the four *Id* genes were induced by BMP4 in NSC cultures (Fig. 3A), only Id1 and Id4 were also upregulated at the protein level (Figs 3B-D and S2B, C). Of those, Id4 expression was highly variable and was not seen in cells presenting high levels of Ascl1, suggesting a possible negative regulatory relationship between the two proteins, while Id1 was also expressed at various levels but without apparent relationship with Ascl1 levels (Figs 3C-E and S2D-I). Co-immunoprecipitation experiments in BMP-treated NSCs confirmed that Id4 binds the E protein E47 and can therefore sequester this factor away from its binding partner Ascl1 (Figs 3F and S2J).

**Figure 3.**
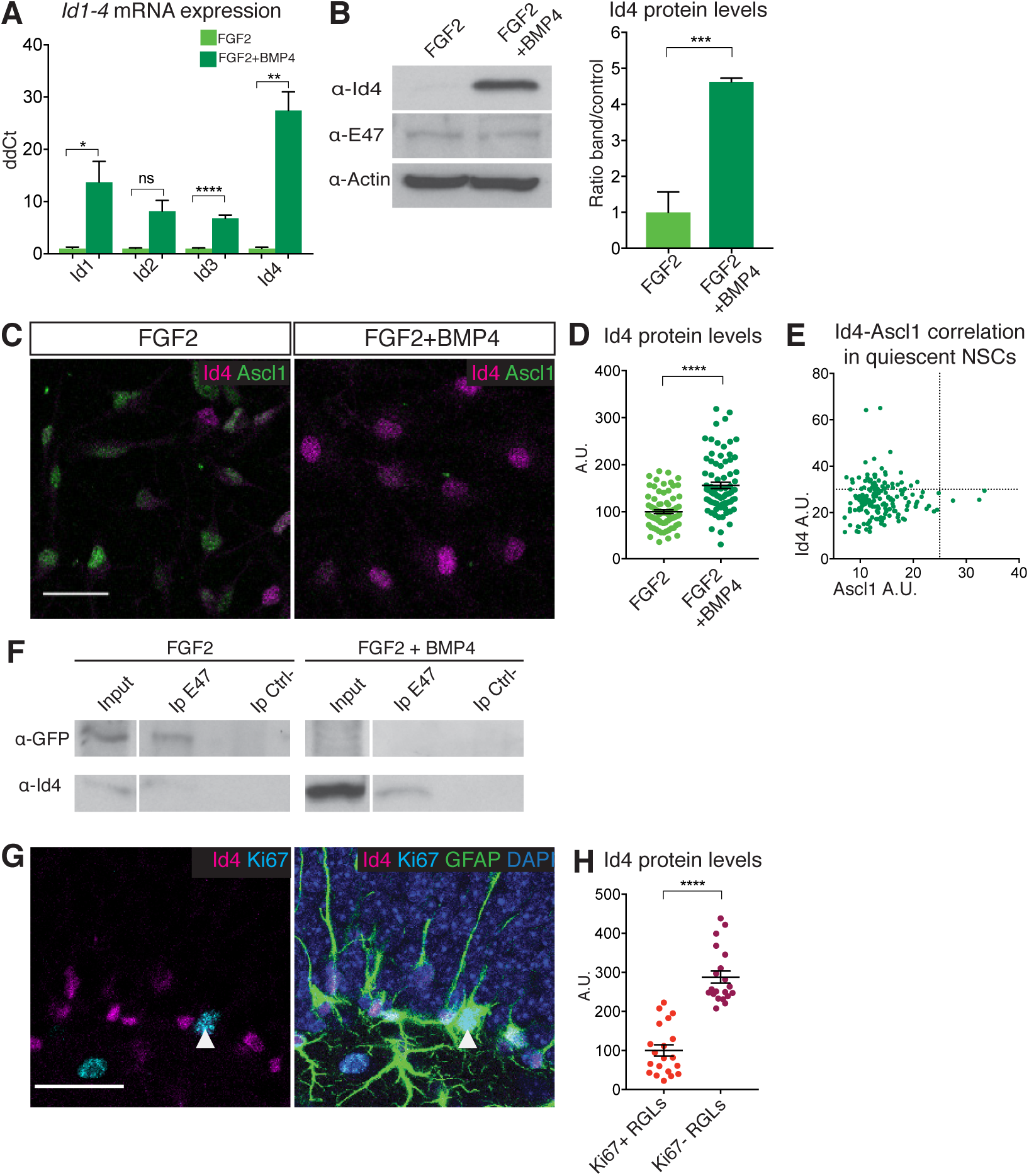
Id4 is a candidate regulator of Ascl1 protein expression in quiescent hippocampal stem cells. (A) Transcript levels for the four Id genes (*Id1*, *Id2*, *Id3*, *Id4*) in FGF2- and FGF2+BMP4-treated NSCs cultures analysed by QPCR. BMP strongly induces the *Id* genes. p>0.05 (ns), p<0.05 (*), p<0.01 (**), p<0.0001 (****). n=3. (B) Western blot analysis and quantification of Id4 and E47 in FGF2- and FGF2+BMP4-treated NSCs. BMP4 upregulates Id4 protein expression; E47 expression is unchanged. p<0.001 (***). n=3 (C) Immunolabeling for Id4 and Ascl1 in FGF2- and FGF2+BMP4-treated NSCs. Scale bar, 30μm. (D-E) Quantifications of the data in (C). (D) BMP4 treatment increases Id4 protein levels in NSCs. p<0.0001 (****). n=3. (E) Id4 protein is expressed at high levels in NSCs expressing low levels of Ascl1 protein. Dotted lines show the threshold for positive staining for each protein. P<0.05 (*), two-sided Fisher’s exact test. n=3. (F) Immunoprecipitation of the Ascl1 dimerisation partner E47 from FGF2- and FGF2+BMP4-treated NSCs, followed by western blot analysis of GFP(Ascl1Venus) and Id4. E47 co-immunoprecipitates with GFP in FGF2 conditions but with Id4 in FGF2+BMP4 conditions. An antibody against V5 is used for the negative control. (G) Immunolabeling for Id4, Ki67 and GFAP and staining for DAPI in hippocampal RGLs. Scale bar, 30μm. (H) Quantification of the data in (G). Id4 is expressed at high levels in quiescent (Ki67-) RGLs and at low levels or is not expressed in proliferating (Ki67+) RGLs. p<0.0001 (****). n=3. See also Figures S2 and S3.

In vivo, Id1, Id3 and Id4 proteins are expressed in the SGZ of the DG where hippocampal stem cells are located, while Id2 is expressed by granule neurons but not in the SGZ (Figs 3G and S3A-E). Id4 was expressed at high levels in quiescent hippocampal RGLs and at much lower levels in proliferating RGLs (Fig. 3G, H). Id1 was expressed in fewer RGLs and was enriched in proliferating cells, while Id3 was only expressed in very few RGLs (Fig. S3A, E). Id4 is therefore a good candidate to suppress Ascl1 protein expression and promote quiescence in hippocampal stem cells, both in culture and in vivo.

### Id4 promotes the degradation of Ascl1 protein and induces a quiescence-like state in NSCs

To address the role of Id4 in Ascl1 regulation and in hippocampal stem cell quiescence, we first asked whether forcing the expression of Id4 in proliferating NSCs would be sufficient to reduce Ascl1 protein level and induce a quiescent state (Fig. 4D). Id4 expression was low in control proliferating NSCs (Fig. 3B-D) and was strongly increased after transfection with an Id4 expression construct (Fig. 4A). Id4-transfected NSCs maintained Ascl1 mRNA at levels similar to those of control NSCs but showed markedly reduced Ascl1 protein levels (Fig. 4A-C). Moreover, transfection of Id4 resulted in a significant decrease in NSC proliferation (Fig. 4E-F). The effects of Id4 protein on Ascl1 expression and NSC proliferation suggest that induction of Id4 in BMP-treated NSCs contributes to the degradation of Ascl1 protein and the induction of quiescence (Fig. 4D).

**Figure 4.**
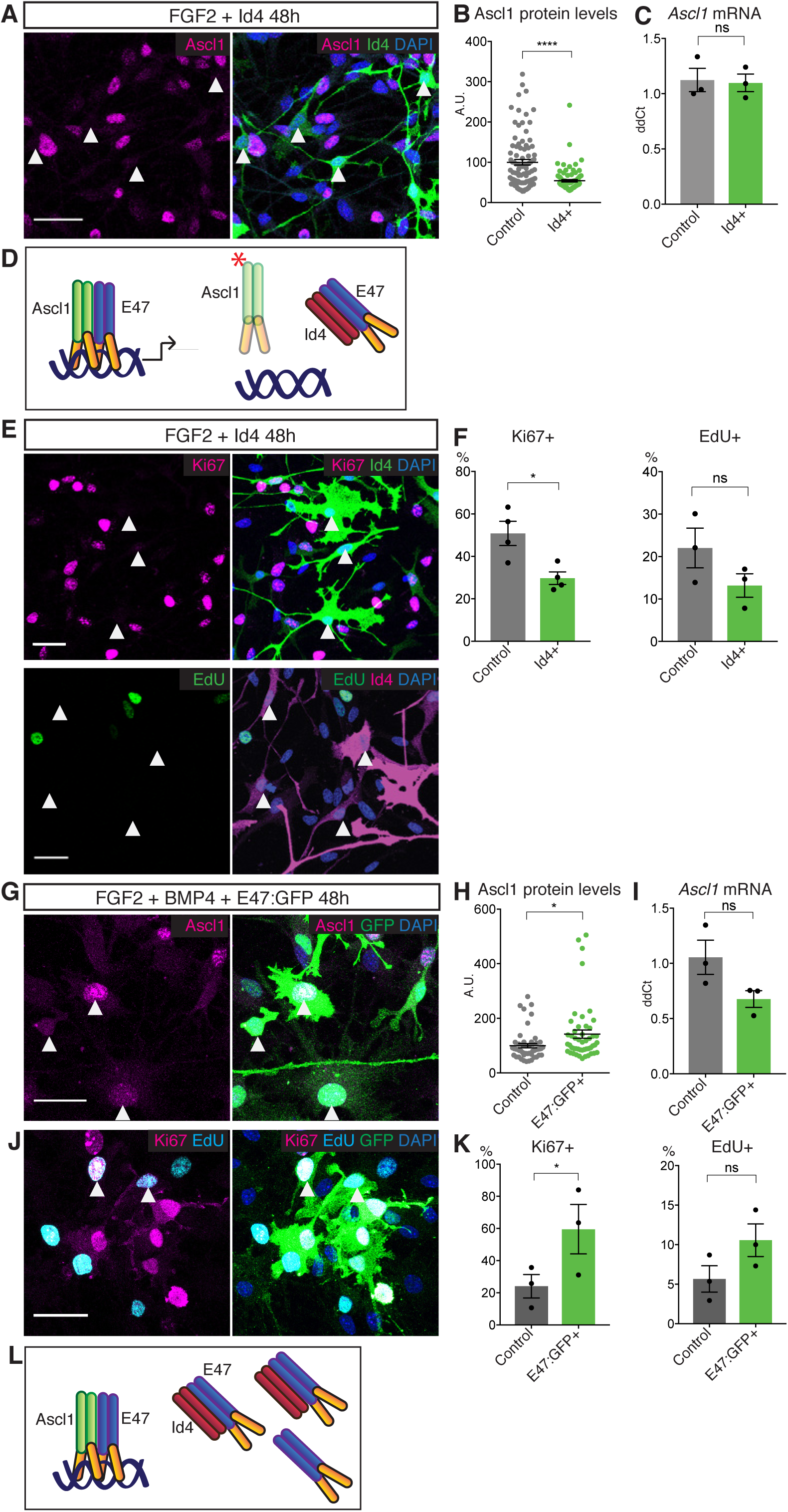
Id4 suppresses Ascl1 expression and cell proliferation in NSC cultures. (A) Immunolabeling for Ascl1 and Id4 and staining for DAPI in Id4-overexpressing, FGF2-treated NSCs. White arrows show low Ascl1 levels in Id4-overexpressing cells. Scale bar, 30μm. (B) Quantification of the data in (A). Ascl1 protein expression is strongly reduced by Id4 overexpression. p<0.0001 (****). The data show one representative experiment, n=3. (C) Ascl1 mRNA levels in FACS sorted FGF2-treated NSCs transfected with a GFP-expressing control or Id4-expression construct. Ascl1 mRNA levels are not changed by Id4 overexpression. p>0.05 (ns). n=3. (D) Model of Ascl1 monomerization and elimination following Id4 overexpression in proliferating NSC cultures. (E) Immunolabeling for Ki67 and Id4 (upper panels) and staining for EdU and immunolabeling for Id4 (lower panels) in Id4-overexpressing, FGF2-treated NSCs. EdU was administered to the cultured cells one hour before fixation. White arrows indicate absence of Ki67 and EdU in Id4-overexpressing cells. Scale bar, 30μm. (F) Quantification of the data in (E). Id4 overexpression reduces NSC proliferation. p>0.05 (ns), P<0.05 (*). n=4 for Ki67 and n=3 for EdU. (G) Immunolabeling for Ascl1 and GFP with DAPI staining, in E47:GFP-overexpressing, FGF2+BMP4-treated NSCs. White arrows indicate Ascl1-positive, E47-overexpressing quiescent cells. Scale bar, 30μm. (H-I) Quantification of the data in (G). Titration of Id proteins by E47 results in a significant increase in Ascl1 protein expression without significant change in Ascl1 RNA levels (I). p>0.05 (ns), P<0.05 (*). n=3 (J) Immunolabeling for GFP and Ki67 and staining for EdU in E47:GFP-overexpressing, FGF2+BMP4-treated NSCs. White arrows indicate Ki67- or EdU-positive, E47-overexpressing quiescent cells. Scale bar, 30μm. (K) Quantification of the data in (J). Titration of Id proteins by E47 reverts the proliferation arrest of BMP4-treated NSCs. p>0.05 (ns), P<0.05 (*). n=3. (L) Model of Id protein titration by E47 overexpression in quiescent NSC cultures.

Next, we asked whether inactivating Id proteins could stabilise Ascl1 protein and revert some aspects of the quiescent state in BMP-treated NSCs. Because NSCs express the four Id proteins, which might have redundant functions, we chose to neutralize all of them by overexpressing the E protein E47. An excess amount of E47 should sequester Id proteins into E47-Id complexes, allowing the formation of Ascl1-E47 complexes and the stabilisation of Ascl1 (Fig. 4L). Indeed, overexpression of E47 in BMP-treated NSCs resulted in an increase in the levels of Ascl1 protein without significantly affecting Ascl1 mRNA levels (Fig. 4G-I). Overexpression of E47 was also sufficient to partially revert the cell cycle arrest of BMP-treated NSCs (Fig. 4J, K). Together, these results support a model whereby induction of high levels of Id4 protein in quiescent NSCs by BMP4 promotes the degradation of Ascl1 by sequestering its dimerization partners (Fig. 4L). They also raise the possibility that suppression of the transcriptional activity of Ascl1 is a key feature of the induction of quiescence by Id4.

### Quiescence is characterised by a downregulation of Ascl1 target genes

To investigate the mechanism by which Id4 induces quiescence in NSCs, we compared the transcriptome of Id4-overexpressing and control proliferating NSCs using RNAseq. Expression of Id4 resulted in the up-regulation of 806 genes and down-regulation of 823 genes (Fig. 5A). Id4-regulated genes represented 44.2% of the genes regulated by BMP4 in NSCs, indicating that Id4 has an important role in the induction of the gene expression programme of quiescence downstream of BMP (Figs 5B and S4A, B). The genes commonly regulated by Id4 and BMP4 are involved in cell cycle (downregulated) and cell adhesion (upregulated) (Fig. 5C-D), which are key hallmarks of the NSC quiescent state (Llorens-Bobadilla et al., 2015; Martynoga et al., 2013; Shin et al., 2015). Direct transcriptional targets of Ascl1 were strongly downregulated in Id4-overexpressing NSCs, including genes with important roles in cell cycle progression such as *Skp2*, *Cdk1*, *Cdk2* and *Foxm1* and other canonical Ascl1 targets such as *Dll1* and *Dll3* (Castro et al., 2011; Martynoga et al., 2013) (Fig. 5E-F).

**Figure 5.**
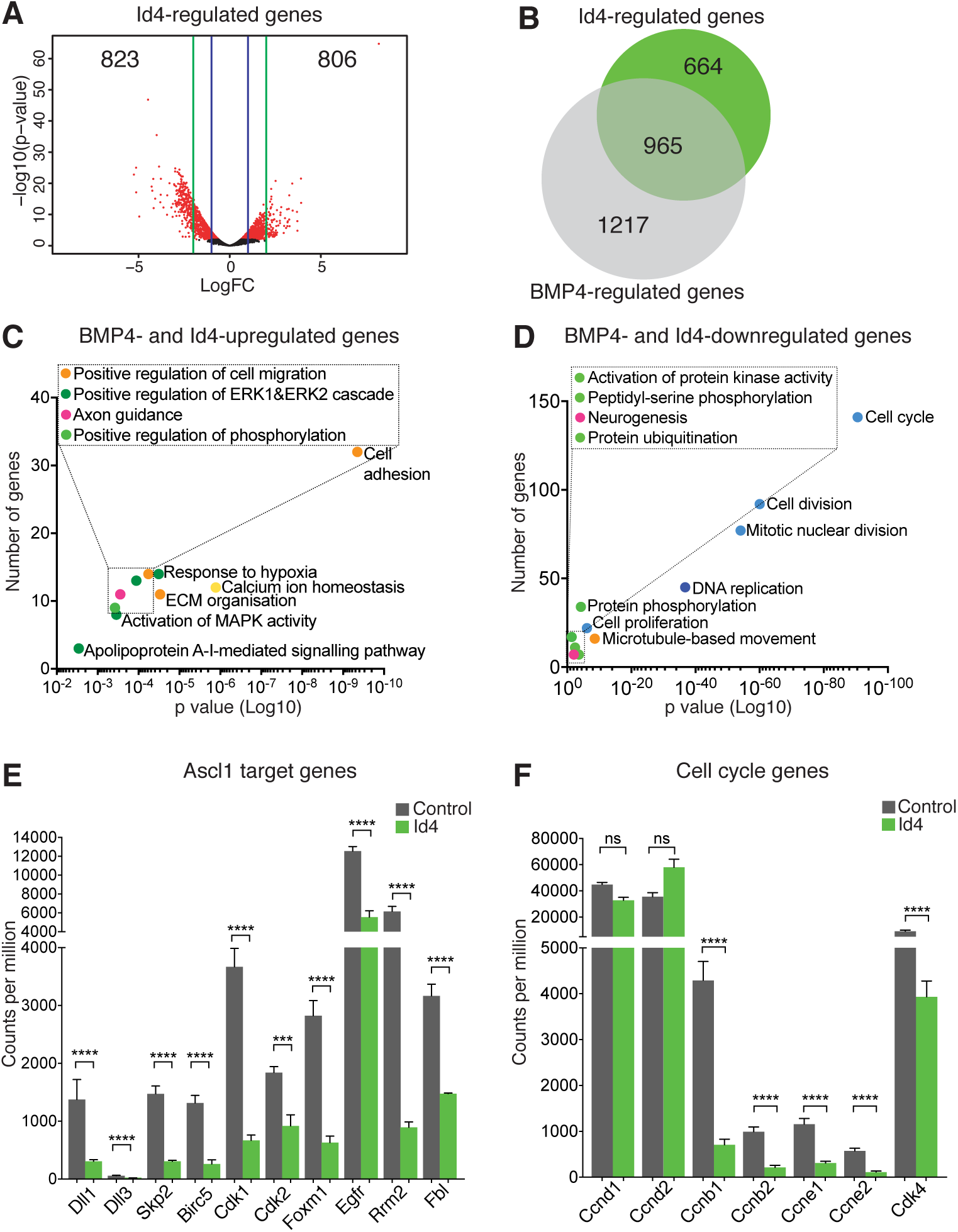
Id4 regulation of Ascl1 targets contributes to BMP-induced NSC quiescence. (A) Volcano plot displaying gene expression changes between control and Id4-overexpressing FGF2-treated NSCs analysed by RNA-Seq. (B) Venn diagram indicating the number of genes de-regulated by addition of BMP4 or Id4 overexpression or both in FGF2-treated cultures. (C-D) Gene Ontology terms associated with genes up- or down-regulated by both addition of BMP4 and Id4 overexpression in FGF2-treated cultures. Dots are coloured based on their ontology terms; light blue: cell cycle/division; dark blue: DNA repair/replication; light green: Protein phosphorylation/modification; dark green: signalling, transcription; orange: adhesion/cytoskeleton; yellow: ion-related; pink: brain/nervous system related. (E) Downregulation of Ascl1 target genes in FGF2-treated cultures overexpressing Id4 and analysed by RNA-Seq, including genes involved in cell cycle regulation (*Skp2*, *Cdk1*, *Cdk2* and *Foxm1*) RGL activation (*Egfr*), canonical Ascl1 targets (*Dll1 and Dll3*) and other Ascl1 targets previously identified in NSCs (*Birc5*, *Rrm2 and Fbl*). p<0.001 (***), p<0.0001 (****). n=3. (F) Down regulation of cell cycle genes in FGF2-treated cultures overexpressing Id4 and analysed by RNA-Seq. p>0.05 (ns), p<0.0001 (****). n=3. See also Figure S4.

We also compared the transcriptome of E47-overexpressing and control BMP-treated NSCs by RNAseq analysis, which revealed the deregulation of 2387 genes. Genes upregulated in E47-overexpressing BMP-treated NSCs represented 40.6% of the genes downregulated when NSCs were exposed to BMP4 (Fig. S4C-E) and included several Ascl1 target genes (Fig. S4F). Overall, our analysis indicates that induction of Id4 by BMP4 and the subsequent degradation of Ascl1 results in the downregulation of its targets, leading to the cell cycle arrest of NSCs (Fig. 4D).

### Loss of Id4 in vivo activates quiescent adult hippocampal RGLs

We next assessed the role of *Id4* in the maintenance of the quiescent state of RGLs in vivo by analysing the hippocampus of mice carrying a conditional mutant allele of *Id4* (*Id4^fl^*) (Best et al., 2014) (Fig. 6A and S5A). To eliminate *Id4* from RGLs, we crossed *Id4^fl/fl^* mice with the Glast-CreERT2 deleter line (Mori et al., 2006) and the Rosa-floxed EYFP reporter line (Srinivas et al., 2001) (Fig. 6A). We administered tamoxifen for 5 days to the triple transgenic mice, which resulted in complete elimination of Id4 protein (Fig. 6B) and we analysed the brains immediately after (*Id4^cKO^* mice; Fig. 6A). The fraction of RGLs expressing Ascl1 increased from 6.0±0.6 in control mice to 15.0±2.1 in *Id4^cKO^* mice (Fig. 6C, D). Ascl1 protein levels in Ascl1-expressing RGLs were also upregulated in *Id4^cKO^* mice (Fig. 6D). The fraction of proliferating RGLs increased from 5.1±1.1 in control mice to 12.3±1.9 in *Id4^cKO^* mice (Fig. 6E, F). When *Id4^cKO^* mice were analysed 30 days after tamoxifen administration and *Id4* deletion, however, the rate of proliferation of RGLs was not significantly different from those found in control mice (Fig. S5D). Id1 levels were upregulated in Id4cKO mice (Fig. S5E); however, this upregulation is already present at P65 (Fig. S5B) meaning that loss of Id1 does not compensate for the loss of Id4. Together, these findings demonstrate that Id4 expression in hippocampal RGLs is essential for the suppression of Ascl1 protein expression and the maintenance of quiescence, and suggest that in the prolonged absence of Id4, a compensatory mechanism maintains RGLs quiescence.

**Figure 6.**
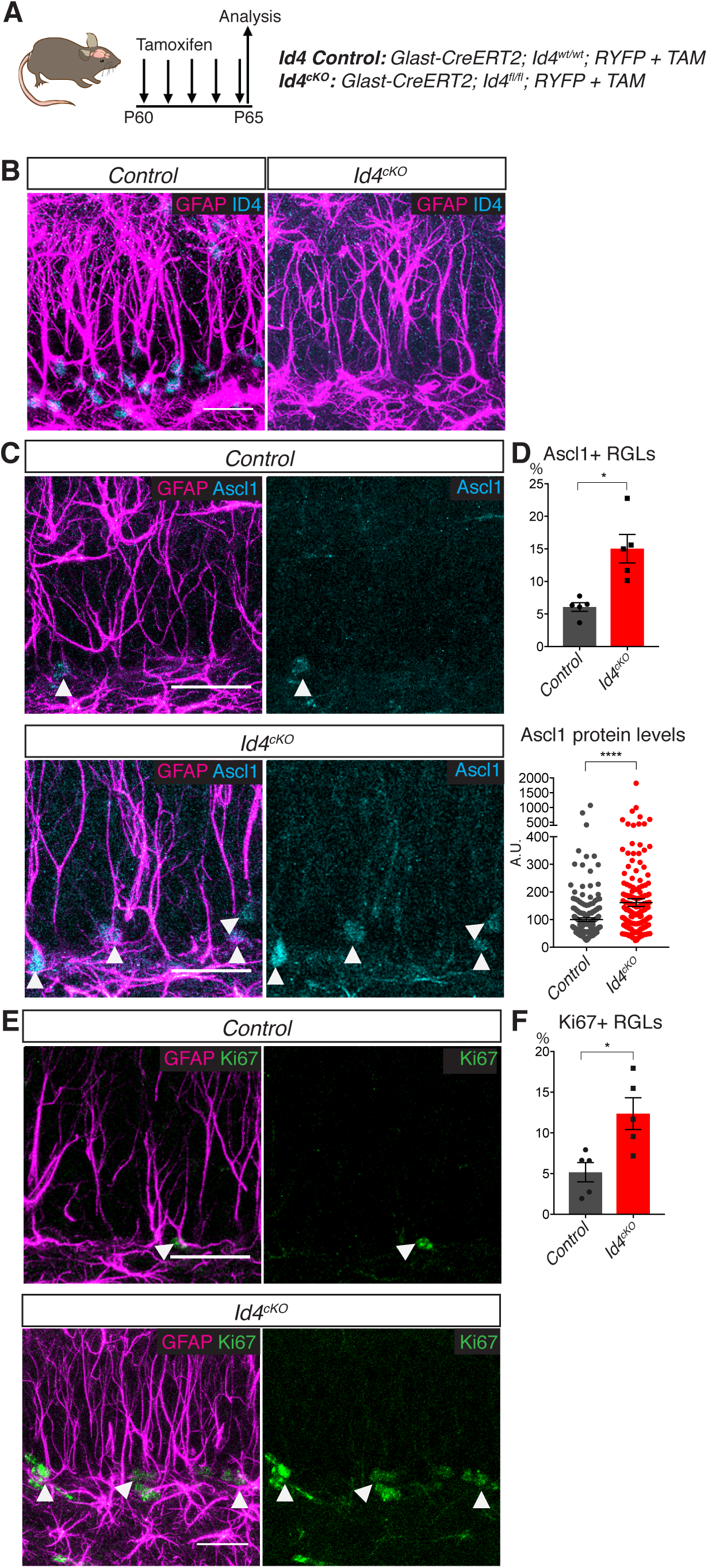
Loss of Id4 results in activation of quiescent RGLs in the adult hippocampus. (A) Design of the experiment of deletion of Id4 from RGLs of the adult hippocampus using *Id4^cKO^* mice. (B) Immunolabeling for GFAP and Id4 demonstrating Id4 elimination in *Id4^cKO^* mice after 5 days of tamoxifen administration. Scale bar, 30μm. (C) Immunolabeling for GFAP and Ascl1 in control and *Id4^cKO^* and control mice after 5 days of tamoxifen administration. White arrows indicate Ascl1-positive RGLs. Scale bar, 30μm. (D) Quantification of the data in (C). Loss of Id4 results in increases in the number of Ascl1-expressing cells and in the levels of Ascl1 protein in RGLs. p<0.05 (*), p<0.0001 (****). n=5 for both control and *Id4^cKO^*. (E) Immunolabeling for GFAP and Ki67 in control and *Id4^cKO^* mice. White arrows indicate Ki67-positive RGLs. Scale bar, 30μm. (F) Quantification of the data in (E). Loss of Id4 results in an increase in the fraction of proliferating RGLs. p<0.05 (*). n=5 for both control and *Id4^cKO^*. See also Figure S5.

### Id4 is regulated by multiple signalling pathways in the adult dentate gyrus

As Id4 function is essential for the maintenance of quiescence of hippocampal RGLs, we examined the signaling pathways that induce expression of the *Id4* gene in these cells. Since BMP4 induces *Id4* in hippocampal NSC cultures (Fig. 3A, B), we asked whether BMP signaling is required for the maintenance of Id4 expression in hippocampal RGLs. We measured Id4 expression in mice in which Smad4, an essential component of the BMP signalling pathway, has been eliminated from hippocampal RGLs (Chu et al., 2004) (Fig. 7A). Unexpectedly, Id4 levels were increased in RGLs from *Smad4^cKO^* mice, while Id1 expression in RGLs was severely reduced in the absence of *Smad4* (Figs 7B-E and S6A-C). Ascl1 expression and RGL proliferation remained similar in *Smad4cKO* and control mice, supporting the idea that *Id4* and not *Id1* is essential to suppress Ascl1 and maintain RGL quiescence (Fig. 7F). As Notch signaling has also been implicated in the regulation of *Id* genes (Liu and Harland, 2003), we examined Id proteins in mice in which *RBPJk*, an essential component of the Notch pathway, has been deleted (GlastCreERT2; *RBPJk^fl/fl^*; RYFP or *RBPJk^cKO^* mice (Han et al., 2002); Fig. 7G). The percentage of Id4+ RGLs was only slightly lower in these mice (Fig. 7H, I), while the fractions of Ascl1+ and proliferating RGLs were elevated compared with control mice, as previously reported (Andersen et al., 2014; Magnusson et al., 2014); Fig. S6D-E), presumably due to transcriptional upregulation of the *Ascl1* gene. We then asked whether the BMP and Notch pathways might act redundantly to maintain Id4 expression, by generating mice in which both *Smad4* and *RBPJk* were deleted (*Smad4^cKO^*;*RBPJk^cKO^* mice; Fig. 7G). Id4 was expressed in a comparable fraction of RGLs in these mice as in control mice, albeit at a slightly reduced level (Fig. 7J, K). Therefore, multiple signalling pathways present in the hippocampal niche, including Notch-RBPJk, BMP-Smad4 and additional unidentified pathways, converge on the induction of Id4 to promote the degradation of Ascl1 and maintain NSC quiescence.

**Figure 7.**
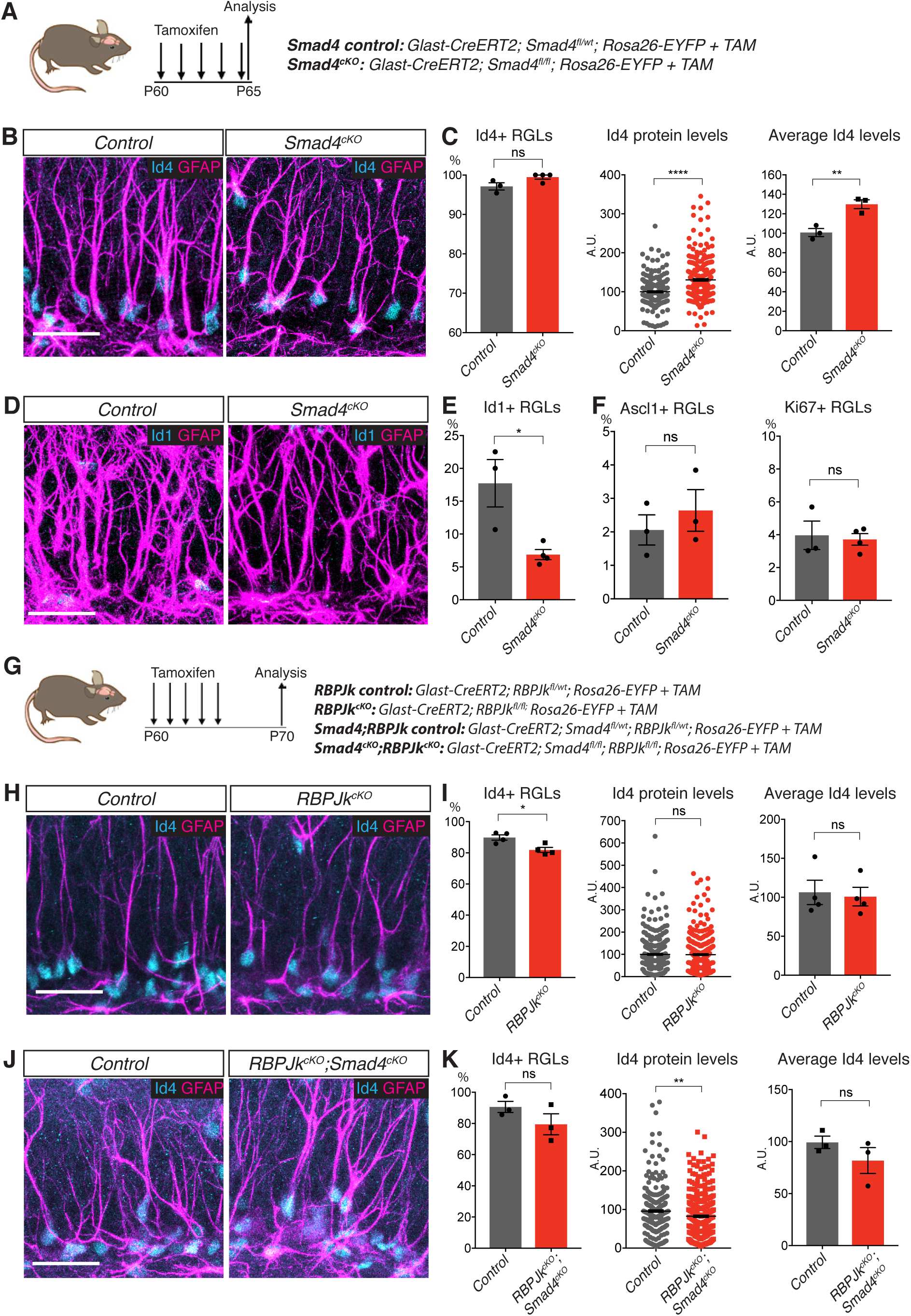
Roles of the Notch and BMP pathways in regulation of Id4 expression. (A) Design of the experiments of inactivation of the BMP pathway in hippocampal RGLs. (B) Immunolabeling for Id4 and GFAP in the DG of *Smad4^cKO^* and control mice 5 days after the first tamoxifen administration. Scale bar, 30μm. (C) Quantification of the data in (B). The number of Id4 expressing RGLs is unchanged while the levels of Id4 are significantly increased in the absence of BMP-Smad4 signaling. p>0.05 (ns), p<0.01 (**), p<0.0001 (****). n=3 for control and n=4 for *Smad4^cKO^*. (D) Immunolabeling for Id1 and GFAP in the DG of *Smad4^cKO^* and control mice 5 days after the first tamoxifen administration. Scale bar, 30μm. (E) Quantification of the data in (D). Id1 expression in hippocampal RGLs is strongly reduced in the absence of BMP-Smad4 signaling. p<0.05 (*). n=3 for control and *Smad4^cKO^*. (F) The fractions of RGLs expressing Ascl1 and proliferating are unchanged in *Smad4^cKO^* mice indicating that a reduction in Id1 does not increase Ascl1 levels or RGL activation. p>0.05 (ns). n=3. (G) Design of the experiments of inactivation of the Notch or Notch+BMP pathways in hippocampal RGLs. More time was allowed after tamoxifen than in (A) to ensure the elimination of the very stable RBPJk protein. (H) Immunolabeling for Id4 and GFAP in the DG of *RBPJk^cKO^* and control mice 10 days after the first tamoxifen administration. Scale bar, 30μm. (I) Quantification of the data in (H). A slightly lower number of RGLs are positive for Id4, while its levels are unaffected by the loss of *RBPJk*. p>0.05 (ns), p<0.05 (*). n=4 for control and *RBPJk^cKO^*. Scale bar, 30μm. (J) Immunolabeling for Id4 in the DG of *Smad4^cKO^*;*RBPJk^cKO^* and control mice 10 days after the first tamoxifen administration. Scale bar, 30μm. (K) Quantification of the data in (J). Expression of Id4 is only moderately reduced in the absence of both *Smad4* and *RBPJk*. p>0.05 (ns), p<0.01 (**). n=3 for control and *Smad4^cKO^*;*RBPJk^cKO^*. See also Figure S6.

## Discussion

In this study, we show that the repressor protein Id4 is essential to maintain adult hippocampal stem cells (RGLs) in a quiescent state. The function of Id4 in maintenance of RGL quiescence is in remarkable contrast with its role in promoting the proliferation of progenitor cells in the embryonic cerebral cortex (Bedford et al., 2005; Yun et al., 2004). This difference may reflect the different role of the bHLH proteins that are Id4 targets in embryonic versus adult neural lineages. In the embryonic forebrain, neural stem cells are in a proliferative state and bHLH proteins mostly act by promoting neuronal differentiation. Their inactivation by Id4 during development therefore results in a block of differentiation and extended proliferation. In contrast, in the adult hippocampus RGLs are mostly quiescent, Ascl1 is required to promote their activity and inactivation of Ascl1 by Id4 results in their failure to proliferate.

We also show that Id4 promotes the degradation of the pro-activation factor Ascl1 in RGLs. As Ascl1 protein is only detectable in a small fraction of proliferating RGLs (Andersen et al., 2014), we were not expecting the Ascl1 gene to be transcribed by most RGLs including many quiescent cells. We found that despite Ascl1 mRNA being expressed and translated, Ascl1 protein does not accumulate in quiescent RGLs due to rapid degradation. This surprising finding could be the reason why single cell transcriptomic analysis of hippocampal cells did not identify Ascl1 among the genes differentially expressed between quiescent and active stem cells (Artegiani et al., 2017; Hochgerner et al., 2018; Shin et al., 2015). This non-transcriptional control of a key activation factor is also found, for instance, in satellite stem cells where the bHLH factor MyoD is transcribed in quiescent cells but its translation is inhibited by an RNA-binding protein to prevent stem cell activation (de Morree et al., 2017).

It is well established that Id proteins, including Id4, form non-functional heterodimers with E proteins, which are dimerization partners of tissue-specific bHLH transcription factors such as Ascl1 (Imayoshi and Kageyama, 2014; Ling et al., 2014; Patel et al., 2015; Sharma et al., 2015). High levels of Ids are therefore expected to result in the sequestration of E-proteins away from functional dimers with Ascl1. Non-dimerised Ascl1 is not able to bind DNA, and this alone could explain why Ascl1 target genes are downregulated in NSCs upon Id4 overexpression or BMP treatment. But how does Id4 prevent Ascl1 protein accumulation? Exposure of different cell types to BMPs has been shown to trigger the proteolytic degradation of Ascl1 (Shou et al., 1999; Vinals et al., 2004). In lung carcinoma cells, the formation of heterodimers with E47 stabilises Ascl1, and induction of Id1 by BMP2 sequesters E47, resulting in degradation of the unstable monomeric form of Ascl1 (Vinals et al., 2004). We propose that a similar mechanism promotes the degradation of Ascl1 when quiescent hippocampal stem cells express high levels of Id4.

Id4 has been proposed to act differently from other members of the Id family (Boareto et al., 2017; Patel et al., 2015; Sharma et al., 2015). Accordingly, the function that we have identified for Id4 in the hippocampus is also distinct from the role reported for other Id factors in the adult ventricular-subventricular zone (V-SVZ), where Id1 and Id3 promote stem cell self-renewal (Nam and Benezra, 2009) and Id1-3 maintain stem cell function by keeping stem cells adherent to their niche environment (Niola et al., 2012). Besides regulating the activity of tissue-specific bHLH transcriptional activators such as Ascl1, Id proteins also interact with bHLH transcriptional repressors of the Hes family (Bai et al., 2007). Direct interaction of Id2 with Hes1 blocks the autorepressive activity of Hes1 protein resulting in its stable expression at a high level. Therefore, Id proteins promote a switch of the expression pattern of Hes proteins from oscillatory, resulting in oscillatory expression of target genes such as Ascl1, to stably high, resulting in constant repression of these targets (Bai et al., 2007; Boareto et al., 2017). Id4 can inhibit the action of Id1-3 proteins by interacting with stronger affinity with them than with the E-proteins (Patel et al., 2015; Sharma et al., 2015). This non-canonical role of Id4 has been proposed to explain how Id4 promotes the proliferation of embryonic neural progenitors (Bedford et al., 2005; Yun et al., 2004) by blocking the interaction of Id1-3 with Hes proteins and thus promoting Hes protein oscillations and progenitor proliferation (Boareto et al., 2017). *Ascl1* is transcribed in most quiescent RGLs, indicating that Hes proteins do not fully repress transcription of *Ascl1* in these cells. However, *Ascl1* expression is strongly enhanced when the Notch effector *gene* Rbpjk is deleted (Fig. S6D), suggesting that the Notch-RBPJk-Hes pathway partially represses *Ascl1* and maintains *Ascl1* expression at low levels. A possible scenario is that Id proteins partially stabilise Hes protein expression and thereby maintain a mild repression of *Ascl1* transcription while Id4, in addition, degrades the remaining low levels of Ascl1 protein to maintain the quiescent state. We provide evidence that Id4 drives Ascl1 degradation but if and how Id4 also interacts with other Id proteins to regulate hippocampal stem cell activity remains to be addressed.

Id4 is not the only Id protein expressed in hippocampal stem cells, as Id1 is also expressed in about 50% of RGLs and all Id proteins are expressed in cultured NSCs. Nevertheless, it is clear that Id4’s role in regulation of stem cell quiescence is unique. While Id4 expression in vivo and in culture is restricted to stem cells that are quiescent and express low levels or no Ascl1 protein, Id1 protein is found in stem cells that are active and express high levels of Ascl1, suggesting that contrary to Id4, it does not promote stem cell quiescence or the degradation of Ascl1. In agreement with this, Id1 has recently been shown to have a role in the activation of hematopoietic stem cells upon stress signals (Singh et al., 2018). Loss-of-function analysis provides direct evidence of the exclusive role of Id4 in the regulation of RGL activity. Deletion of *Id4* leads to a rapid increase in Ascl1 expression and in the fraction of RGLs that proliferate. The effect of loss of *Id1* can be examined in *Smad4cKO* mice where Id1 expression in RGLs is greatly reduced while Id4 expression is unaffected. *Smad4cKO* do not show defects in Ascl1 expression or RGL proliferation. Why Id1, which has been shown to dimerise with E proteins and promote Ascl1 degradation in another cell type (Vinals et al., 2004) has no such effect in hippocampal stem cells is unclear. It is unknown whether Id4 also has a role distinct from that of other Id proteins in other adult stem cell populations as well.

While loss of Id4 results after a few days in a premature activation of quiescent RGLs, RGL activity and Ascl1 expression are back to normal levels after 30 days. Thus, other Ids or unrelated factors might be induced by the loss of Id4 and compensate for it, as for instance Id1 is (Fig S5B, E). We have previously shown that the ubiquitin ligase Huwe1 degrades Ascl1 in proliferating RGLs and allows a fraction of these cells to return to quiescence (Urban et al., 2016). Huwe1 function is not required in quiescent RGLs since loss of *Huwe1* does not affect the rate at which quiescent RGLs become active. Huwe1 is expressed in quiescent RGLs (Urban et al., 2016) and although Id4 might mask its role in degrading Ascl1 in these cells, it might be able, in the absence of Id4, to eliminate excess Ascl1 and maintain RGL quiescence.

Our transcriptomic analysis suggests that collectively, Id proteins contribute to a large extent to the induction by BMP of a gene expression programme promoting NSC quiescence. Overexpression of E47 in BMP-treated NSCs results in down regulation of a large fraction of the genes that are induced by BMP4 in NSCs, and in induction of a large fraction of the genes suppressed by BMP4. Conversely, overexpression of Id4 in the absence of BMP induces a large fraction of the genes that BMP4 induces and suppresses a large fraction of the genes suppressed by BMP4. However a subset of genes are induced or suppressed by BMP4 independently of Id4. Gene Ontology analysis of the regulated genes indicates that Id4 acts downstream of BMP signaling to promote multiple aspects of NSC quiescence including the induction of cell adhesion molecules and the downregulation of cell cycle genes, many of which are Ascl1 targets.

Given the important role of Id4 in maintaining RGL quiescence, it seems likely that its expression is regulated by niche signals to control RGL activity. Id genes, including Id4, are well known effectors of BMP signaling in neural cells and other cell types (Ling et al., 2014; Patel et al., 2015; Samanta and Kessler, 2004). While Id4 expression is strongly induced by BMP4 in cultured NSCs, deletion of the BMP signaling effector Smad4 in RGLs does not reduce its expression and in fact significantly upregulates it, possibly due to a reduction in Id1 expression and cross-regulatory interactions between Id1 and Id4. The maintenance of Id4 expression in RGLs lacking BMP-Smad4 signaling suggests that another signaling pathway regulates Id4 in these cells. TGFß signaling activity in embryonic midbrain progenitors is mediated in a redundant manner by Smad4 and the transcriptional co-regulator Trim33 (Falk et al., 2014). Whether Trim33 is also required to induce Id4 in hippocampal RGLs downstream of BMP or whether another niche signal contributes to Id4 regulation, remains to be determined. Nevertheless, the reduction of Id1 expression from RGLs in *Smad4cKO* mice indicates that Id4 diverges from other Id proteins not only in its activity but also in the regulation of its expression. Id4 has been shown to be directly regulated by Notch signalling in embryonic neural progenitors (Li et al., 2012) and in adult hippocampal stem cells (Zhang et al., 2018) but we find that Id4 expression is only slightly downregulated in RGLs lacking the essential Notch signaling effector RBPJk. The downregulation of Id4 in adult hippocampal stem cells lacking the Notch2 receptor is however observed only transiently at 2 days after Notch2 deletion and not 19 days later (Zhang et al., 2018), consistent with the idea that other niche-induced pathways can compensate for the lack of Notch signaling and activate Id4 expression. Id4 expression is only mildly affected in RGLs lacking both Smad4 and RBPJk, indicating that additional pathways beside BMP-Smad4 and Notch-Rbpjk promote RGL quiescence by maintaining Id4 expression.

Id4 is expressed in most quiescent RGLs but it is sharply downregulated in active RGLs. Indeed, it is one of the most differentially expressed genes in quiescent versus active stem cells, both in vivo and in NSC cultures (Shin et al., 2015). Down-regulation of Id4 is likely to be crucial for RGLs to leave the quiescent state and become active, and our results suggest that multiple pathways converge on the regulation of Id4 expression, emphasising the importance of this gene in the maintenance of RGL quiescence.

## STAR*METHODS

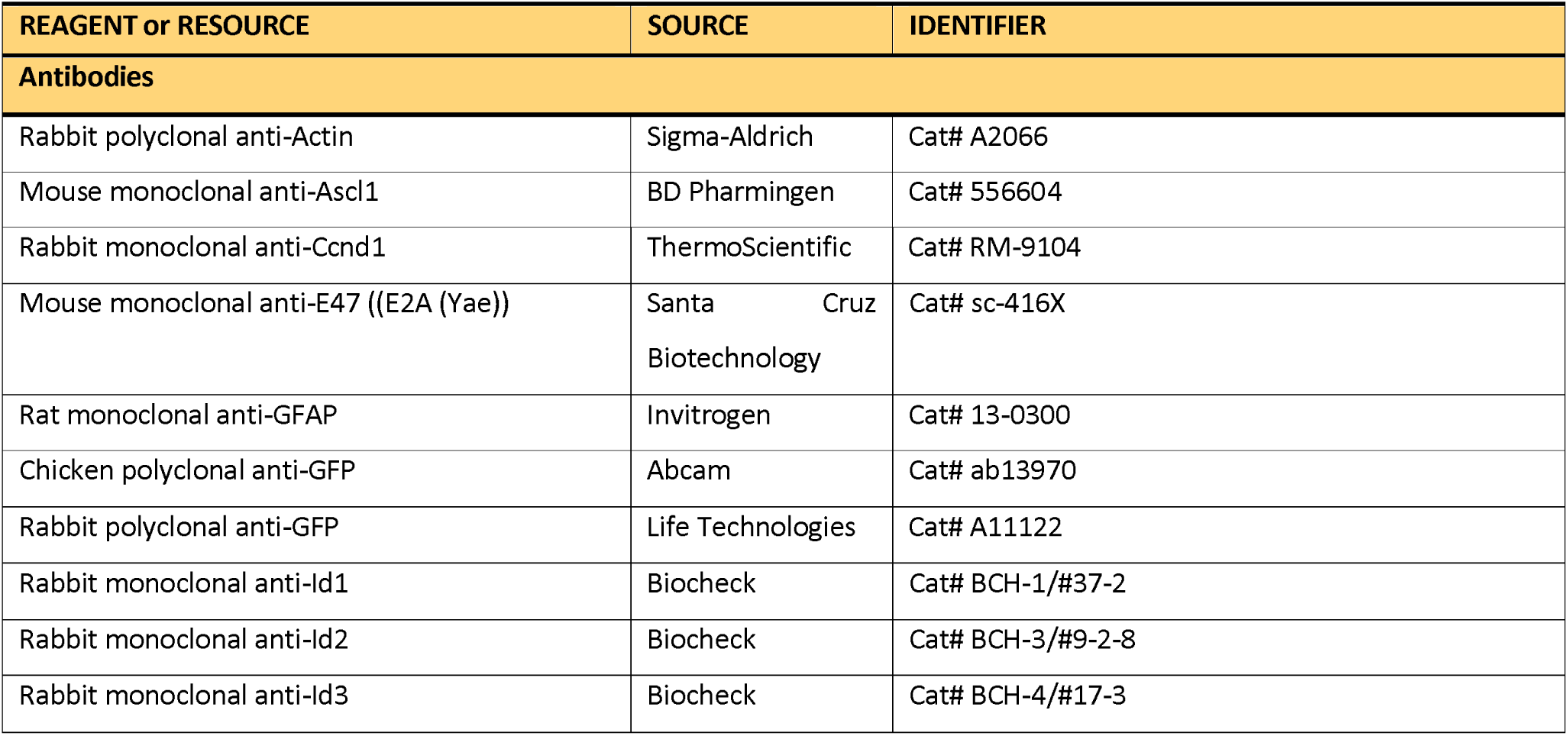
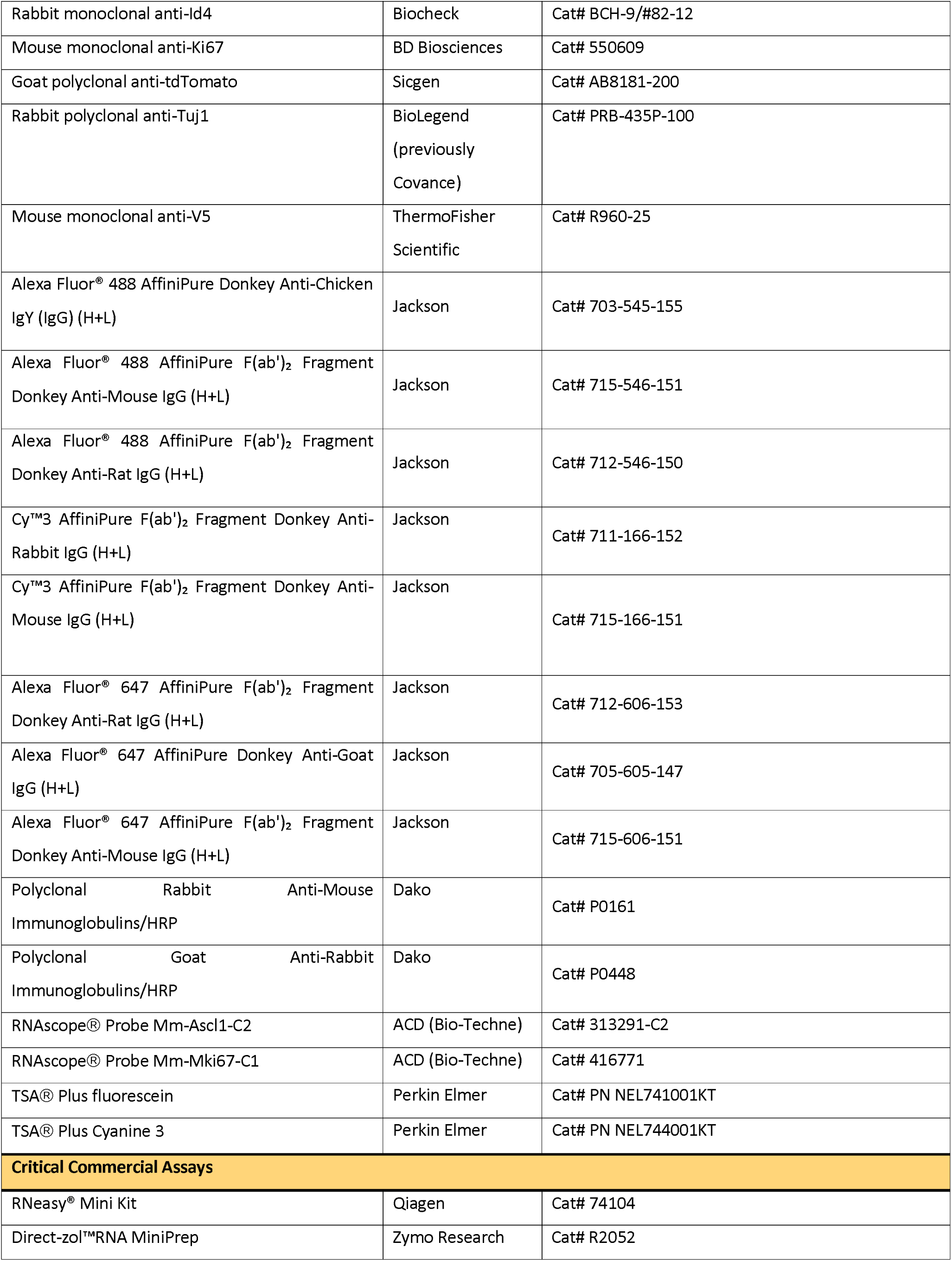
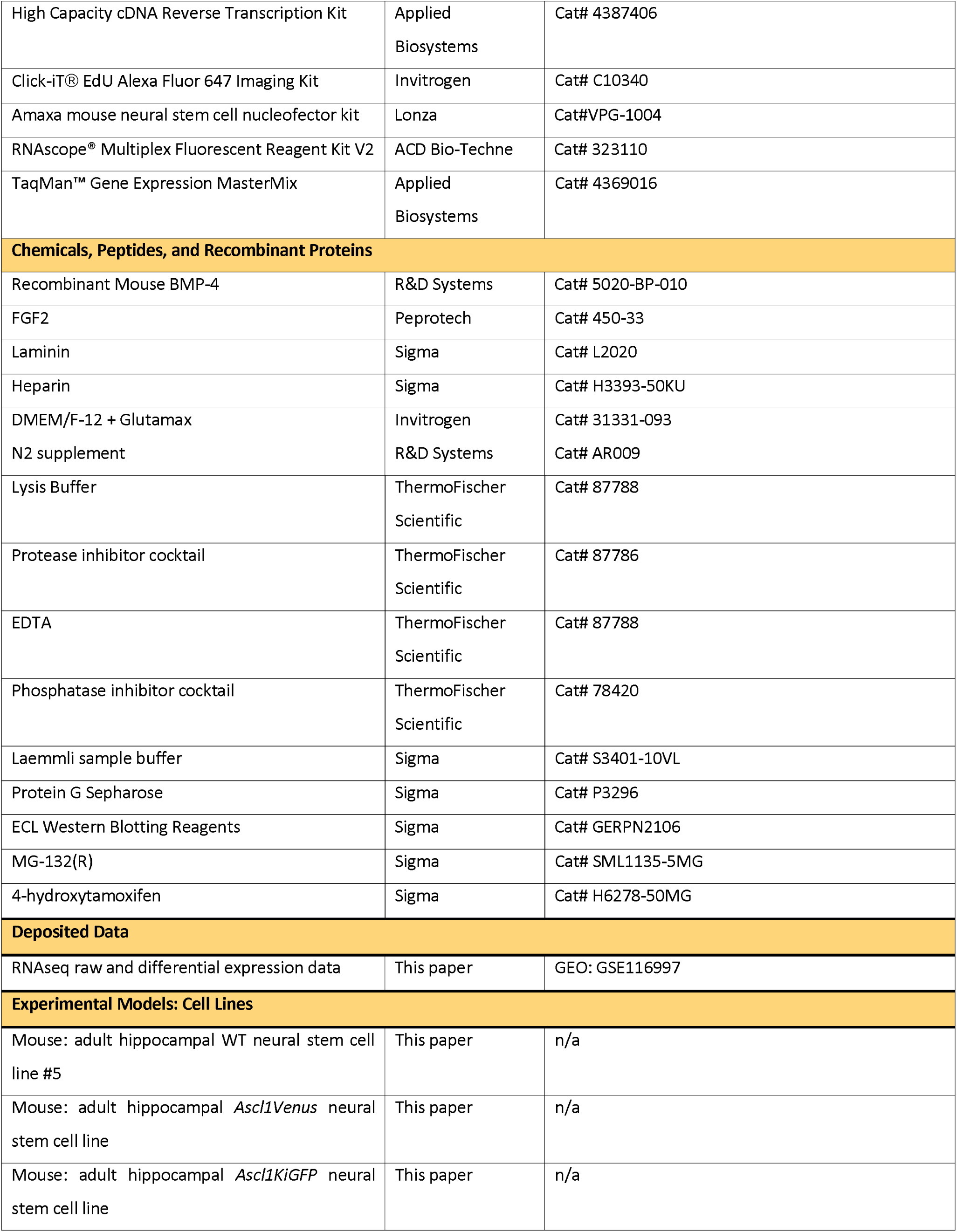
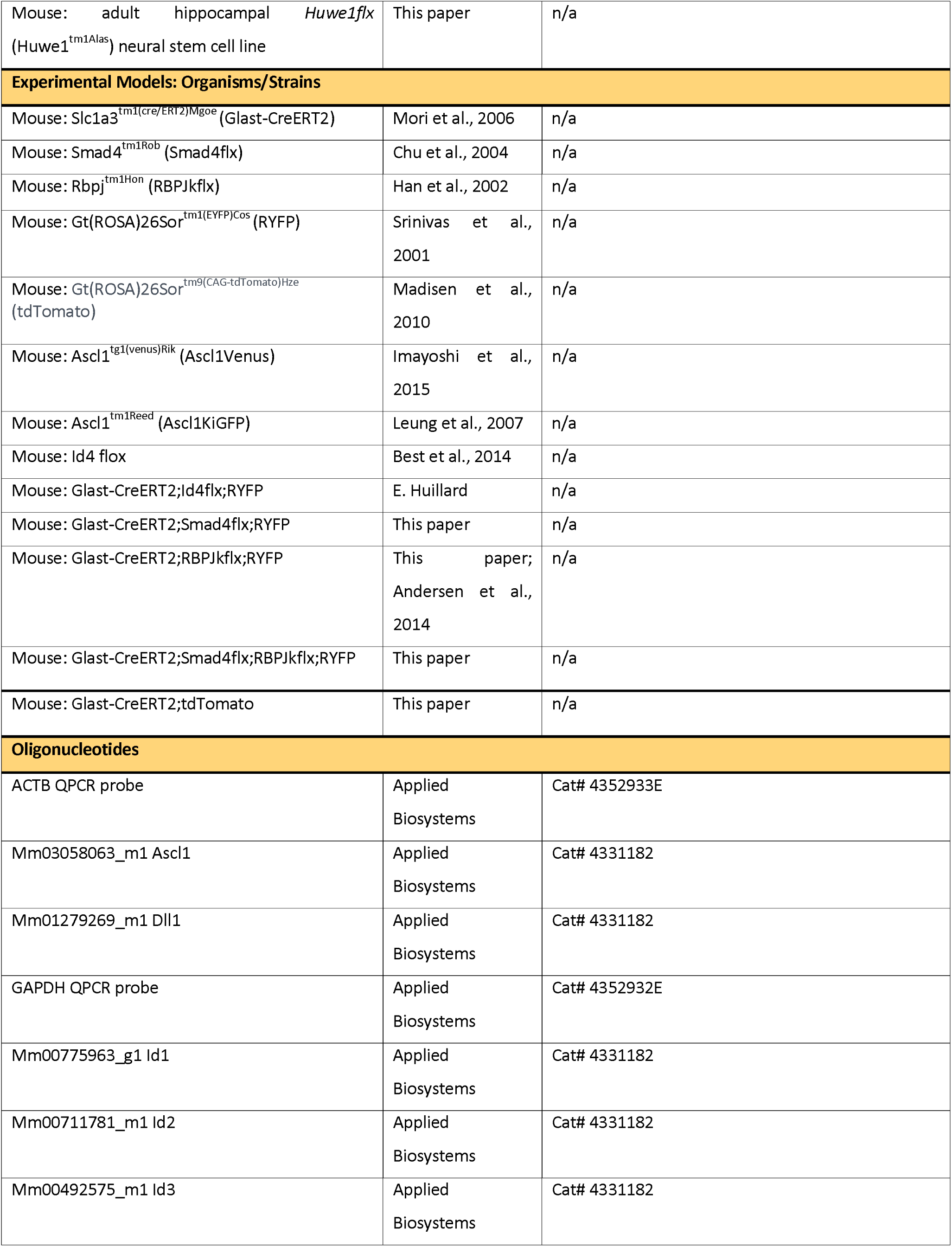
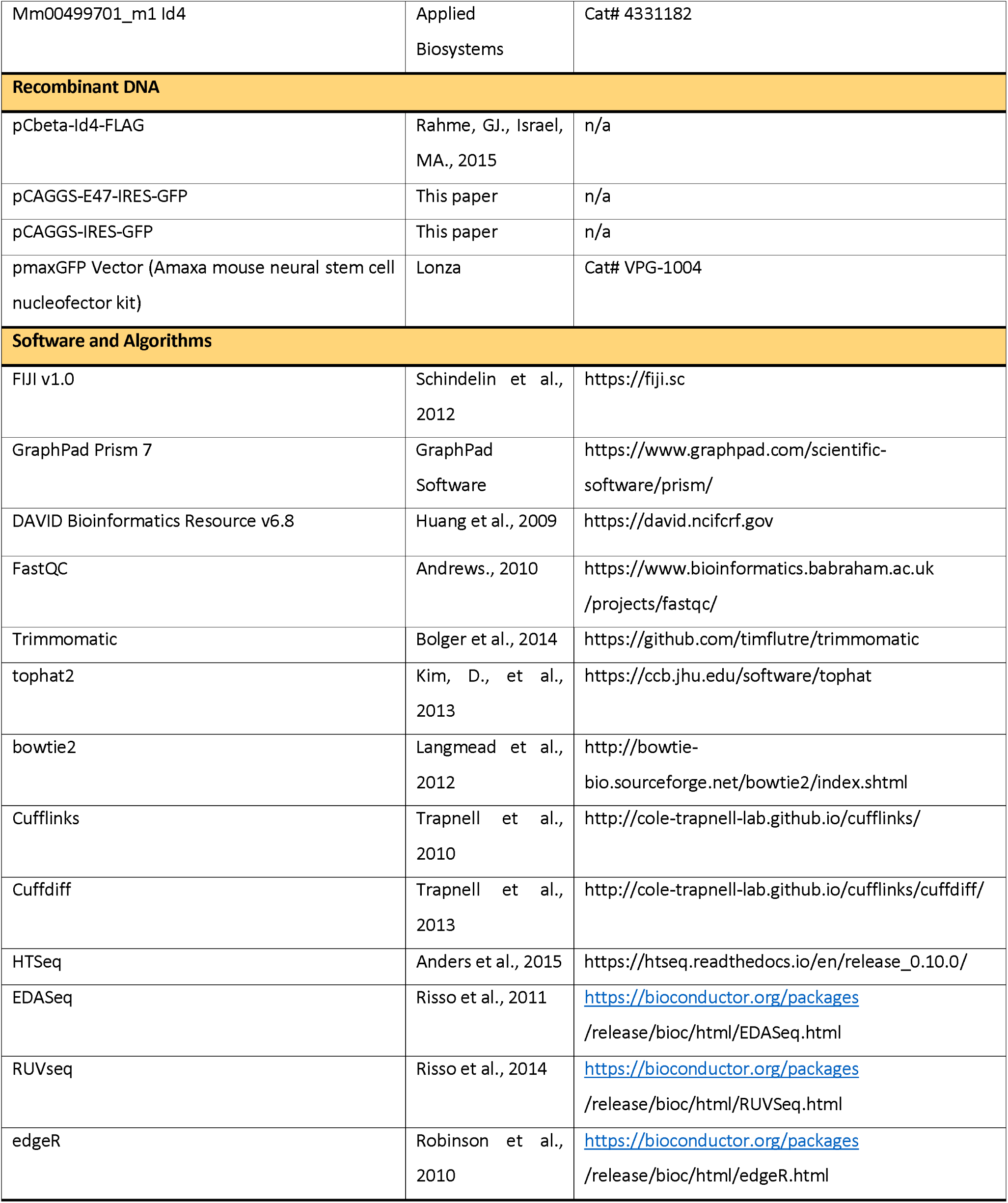

### Experimental model and subject details

#### Mouse models

All procedures involving animals and their care were performed in accordance with the guidelines of the Francis Crick Institute, national guidelines and laws. This study was approved by the Animal Ethics Committee and by the UK Home Office (PPL PB04755CC). Mice were housed in standard cages under a 12h light/dark cycle, with ad libitum access to food and water.

All experimental mice were of a mixed genetic background. Founder mice were bred to MF1 mice, and then backcrossed to littermates of the F1 generation. In order to generate mice with a hippocampal stem cell-specific, tamoxifen-inducible recombination, plus a YFP reporter of recombination, GLAST-CreERT2 (Slc1a3^tm1(cre/ERT2)Mgoe^) (Mori et al., 2006) mice were crossed with Rosa26-floxed-stop-YFP (RYFP; Gt(ROSA)26Sor^tm1(EYFP)Cos^) (Srinivas et al., 2001) mice. These mice were then crossed with our experimental strains:

Smad4flx (Smad4^tm1Rob^) mice, originally reported by (Chu et al., 2004).

RBPJkflx (Rbpj^tm1Hon^) mice, originally reported by (Han et al., 2002).

Ascl1Venus (Ascl1^tg1(venus)Rik^) mice, originally reported by (Imayoshi et al., 2013).

Ascl1KiGFP (Ascl1^tm1Reed^) mice, originally reported by (Leung et al., 2007).

Id4flx mice, originally reported by (Best et al., 2014).

Glast-CreERT2 mice were crossed with Gt(ROSA)26Sor^tm9(CAG-tdTomato)Hze^ (tdTomato), originally
reported by (Madisen et al., 2010).

Glast-CreERT2;Smad4flx;RYFP mice were crossed with Glast-CreERT2;RBPJk;RYFP mice in order to generate the quadruple transgenic Glast-CreERT2;Smad4flx;RBPJkflx;RYFP line.

Both male and female mice were used for all *in vivo* genetic studies. Experimental groups were a mix of animals from different litters for each particular strain. All mice were injected with tamoxifen at postnatal day 60 +/- 2, and brain tissue collected by transcardial perfusion at 2, 5, 10 or 30 days after the first injection.

#### Primary Cell Cultures

For the derivation of adult hippocampal stem cell lines, 7-8 week old mice were sacrificed and the dentate gyrus dissected (previously described by Walker et al., 2013). Cultures were amplified as neurospheres for two passages before dissociation to adherent cultures. Cells were propagated in basal media (DMEM/F-12 + Glutamax (Invitrogen 31331-093) + 1x Neurocult Supplement (Stem Cell Technologies, 05701) + 1x Penicillin-Streptomycin (ThermoFischer Scientific, 15140)+ 2μg/ml Laminin (Sigma, L2020) + 20 ng/ml FGF (Peprotech, 450-33) + 5μg/mL Heparin (Sigma, H3393-50KU). Cells were incubated at 37°C, 5% CO_2_.

The control adult hippocampal neural stem cell line (AHNSC line #5) was derived from a single male WT/RYFP mouse. AHNSC Ascl1Venus cell line was derived from a single male Ascl1^wt/Venus^ mouse. Huwe1 is X-linked, therefore AHNSC Huwe1flx cell line was derived from a male Glast-CreERT2^wt/wt^; Huwe^fl/Y^;Rosa^YFP/YFP^ mouse.

### Method Details

#### Tamoxifen administration

To induce activation of CreERT2 recombinase, 2mg (57-67mg/Kg) of 4-hydroxytamoxifen (Sigma, H6278) was administered intraperitoneally (ip) to mice at postnatal day 60 (P60), at the same time each day for 5 consecutive days. For in situ hybridization experiments, Glast-CreERT2;tdTomato (Ai9) mice received a single injection at postnatal day 60 +/- 2, and brain tissue collected by transcardial perfusion 48h later.

#### Tissue preparation and immunofluorescence

Mice were transcardially perfused with phosphate-buffered saline (PBS) for 3mins, followed by 4% paraformaldehyde (PFA) in PBS for 12mins. Brains were post-fixed for 2hours in 4% PFA at 4°C and washed with PBS. Brains were coronally sectioned at a thickness of 40μm using a vibratome (Leica).

For in situ samples, mice were perfused with PBS for 3mins, followed by 10% neutral buffered formalin (NBF) for 12mins. Brains were post-fixed in 10% NBF at room temperature for 16-32h, and then washed with 70% EtOH. Brains were paraffin embedded, and coronally sectioned at a thickness of 5μm.

Cultured cells were fixed with 4%PFA in PBS for 10mins at room temperature, and washed with PBS.

For immunofluorescence of tissue, samples were blocked with 10% normal donkey serum (NDS) in 1%Triton-PBS for 2hrs at room temperature with rocking. Fixed cells were blocked with 10%NDS in 0.1%Triton-PBS for 1hr at room temperature. Primary antibodies were diluted in 10%NDS in 0.1%Triton-PBS, and incubated with samples overnight at 4°C with rocking. Following 3x 0.1%Triton-PBS washes, samples were incubated with secondary antibodies diluted in 10%NDS in 0.1%Triton-PBS for 2hrs at room temperature with rocking. Following 3x 0.1%Triton-PBS washes, samples were incubated with DAPI 1:10,000 in 1:1 PBS:H_2_O for 30mins at room temperature with rocking. Primary and secondary antibodies are listed in Key Resources Table, dilutions listed in Table S1.

EdU was detected following secondary antibody incubation, using Click-iT^®^ EdU Alexa Fluor 647 Imaging Kit (Invitrogen, C10340), following manufacturer’s instructions.

#### RNA *in situ* hybridization

For RNA *in situ* hybridization, the RNAscope^®^ Multiplex Fluorescent Reagent Kit V2 (ACD Bio-Techne, 323110) was used with NBF fixed-paraffin embedded 5μm sections, and stained according to the standard company protocol. Target retrieval was performed for 15mins, and Protease Plus treatment was carried out for 30mins. For dual RNAscope^®^-immunofluorescence, following the development of HRP-C3 signal and wash steps, slides were washed in distilled H_2_O, and washed 3x 5mins in 0.1%Triton-PBS at room temperature. Slides were then processed for immunofluorescence as described above. Probes and fluorophores are listed in Key Resources Table, dilutions listed in Table S1.

#### Microscopic analysis

All images were acquired using an SP5 confocal microscope (Leica). For cell culture immunofluorescence, 3 random regions of each coverslip were imaged with a z-step of 1μm. For adult tissue immunofluorescence, both left and right dentate gyri of every twelfth 40μm section along the rostrocaudal length of the DG were imaged, with a z-step of 1μm through the whole 40μm section. For quantification of %+ RGLs, at least 200 RGLs in each of at least 3 mice for each genotype were quantified.

RGLs were identified based on their characteristic morphology (nucleus in the subgranular zone, radial process projecting through the molecular layer) and positive labelling with GFAP and GFP in the case of Glast-CreERT2;RYFP recombined cells, or tdTomato positivity in the case of Glast-CreERT2;tdTomato recombined cells.

#### Cell treatments, constructs and transfection

For culturing adult hippocampal NSCs in proliferation conditions, cells were grown in basal media (DMEM/F-12 + Glutamax (Invitrogen, 31331-093)) + 1x N2 supplement (R&D Systems, AR009) + 1x Penicillin-Streptomycin (ThermoFischer Scientific, 15140) + 2μg/ml Laminin (Sigma, L2020) + 5μg/mL Heparin + 20 ng/ml FGF2 (Peprotech, 450-33). To induce quiescence, cells were plated into flasks or onto coverslips in proliferation conditions and allowed to adhere overnight. Media was replaced the next day with basal media or basal media plus 20ng/mL recombinant mouse BMP4 (R&D Systems, 5020-BP), and cultured for 72h at 37°C, 5% CO_2_.

To test that BMP4-induced cells could reactivate and differentiate, BMP4-treated cells were detached from their flask using Accutase (Sigma, A6964) and re-plated into proliferation conditions, and fixed at 24h, 48h and 72h post-reactivation. To differentiate the cells, following 72h reactivation, cells were cultured in the presence of 10ng/mL FGF2 and 2% foetal bovine serum, for 72h.

In order to visualise S-phase, EdU (Invitrogen, C10340) was added to the media of cells in culture 1hr prior to fixation.

To inhibit the proteasome, cells were grown on 10cm diameter dishes for 72h in supplemented basal media with either just 20ng/mL FGF2 or FGF2 + 20ng/mL BMP4. Cells were treated with either 10μM MG132 (Sigma, SML1135) or an equal volume of DMSO (Sigma), for 30, 60 or 120mins.

For overexpression of Id4 and E47 in NSCs, 5×10^6^ cells per construct were nucleofected using the Amaxa mouse neural stem cell nucleofector kit (Lonza, VPG-1004) and Amaxa Nucleofector II (Lonza), using the program A-033, according to manufacturer’s instructions. The pCbeta-Id4-FLAG construct was a kind gift from M. Israel (Rahme et al., 2015). The E47 expression construct was generated by cloning E47 into pCAGGS-IRES-GFP via EcroRV/Xho1. In order to FACS sort Id4-transfected cells, cells were co-transfected with an empty pCAGGS-IRES-GFP construct at half the concentration of pCbeta-Id4-FLAG, to increase the likelihood of GFP+ cells also being Id4+. For FACS and subsequent RNAseq analysis, FGF2 and FGF2+BMP4 control samples were nucleofected with pMaxGFP vector from the Amaxa kit (Lonza, VPG-1004). Following transfection, cells were plated into flasks and onto glass coverslips, in supplemented basal media, and incubated for 48h at 37°C, 5% CO_2_. Cells transfected with Id4 were cultured in the presence of 20ng/mL FGF2; cells transfected with E47 were cultured in the presence of both 20ng/mL FGF2 and 20ng/mL BMP4.

#### FAC sorting

FACS tubes were pre-coated with 5%BSA-PBS at 37°C for at least 30mins prior to sorting. Cells were detached from flasks using Accutase (Sigma) and centrifuged at 0.3RCF for 5mins. Cell pellets were resuspended in 750μL recovery media (5%BSA-PBS + 20 ng/ml FGF + 1μg/mL Heparin). 1μL propidium iodide was added to cell suspensions to check for cell viability. Cells were sorted on a FACS Aria III machine, into recovery media. Both GFP positive and negative cells were recovered into separate tubes.

#### RNA extraction, cDNA synthesis and QPCR

For FACS experiments, cells were lysed using Qiagen lysis buffer. For all other experiments, cells were lysed with Trizol reagent. RNA was extracted using RNeasy^®^ Mini Kit (Qiagen, 74104) or Direct-zol^™^ RNA MiniPrep Kit (Zymo Research, R2052), according to manufacturer’s instructions.

cDNA was synthesised using the High Capacity cDNA Reverse Transcription Kit (Applied Biosystems, 4387406) following manufacturer’s instructions. Gene expression level was measured using TaqMan Gene expression assays (Applied Biosystems) and quantitative real-time PCR carried out on a QuantStudio Real-Time PCR system (ThermoFisher). Gene expression was calculated relative to endogenous controls Gapdh and ActinB, and normalised to the expression of the control sample in each group, to give a ddCt value.

#### RNA sequencing and analysis

RNA concentration was quantified using the Qubit dsDNA BR/HS Assay Kit. A KAPA mRNA HyperPrep Kit (for Illumina) (KAPA Biosystems, Wilmington, MA, USA) was used with 1000ng of RNA diluted to a final volume of 50µl. Each RNA sample was captured with 50µl of capture beads at 65°C for 2 min and 20°C for 5 min. For the second capture, 50µl of RNase free water was used at 70°C for 2 min and 20°C for 5 min. Captured RNA was subjected to the KAPA Hyper Prep assay: end-repair, A-tailing, and ligation by adding 11µl of Fragment, Prime and Elite Buffer (2X). To obtain a distribution of 200-300bp fragment on the library, the reaction was run for 6 min at 94°C. cDNA synthesis was run in 2 steps following manufacturer’s instructions. The ligation step consisted of a final volume of 110 μL of the adaptor ligation reaction mixture with 60μL of input cDNA, 5 μL of diluted adaptor and 45μL of ligation mix (50µL of ligation buffer+ 10 μL of DNA ligase). The Kapa Dual-Indexed Adapters (KAPA Biosystems-KK8720) stock was diluted to 7µM (1.5mM or 7nM) to get the best adaptor concentration for library construction. The ligation cycle was run according to manufacturer’s instructions. To remove short fragments such as adapter dimers, 2 AMPure XP bead clean-ups were done (0.63 SPRI and 0.7SPRI). To amplify the library, 7 PCR cycles were applied to cDNA KAPA HP mix. Amplified libraries were purified using AMPure XP. The quality and fragment size distributions of the purified libraries was assessed by a 2200 TapeStation Instrument (Agilent Technologies, Santa Clara, CA, USA).

Libraries were sequenced with Hiseq4000 (Illumina), 50-bp paired-end reads for sequencing proliferating vs quiescent NSCs; 75bp single-end reads for Id4/E47 overexpressing NSCs, with a depth of 30×10^6^ reads.

The quality of RNA sequence reads was evaluated using FastQC (version 0.11.2)(Andrews, 2010). Low quality reads and contaminants (e.g. sequence adapters) were removed using Trimmomatic (version 0.32) (Bolger et al., 2014). Sequences that passed the quality assessment were aligned to the mm10 genome using tophat2 (version 2.0.14) (Kim et al., 2013), with bowtie2 (version 2.1.0) (Langmead and Salzberg, 2012) or for the quiescent NSC RNAseq data set, Cufflinks (Trapnell et al., 2010). Transcript abundance level (transcript count) was generated using HTSeq (version 0.5.3p9) (Anders et al., 2015). The transcript counts were further processed using R software environment for statistical computing and graphics (version 3.4.0). Data normalization, removal of batch effect and other variant was performed using EDASeq R package [(Risso et al., 2011) and RUVseq package (Remove Unwanted Variation from RNA-Seq package) (Risso et al., 2014). Differential expression was performed using edgeR R package (Robinson et al., 2010), using the negative binomial GLM approach, or for the quiescent NSC RNAseq data set, Cuffdiff (version 7) (Trapnell et al., 2013). Differentially expressed genes with false discovery rate (FDR <=0.05, Benjamini-Hochberg multiple testing correction), expression level in control samples > 1 CPM (counts per million) or >1 FPKM (fragments per kilobase of transcript per million mapped reads) for the quiescent NSC RNAseq data set, and log fold change >1 were retained and used for further processing, gene ontology and pathway analysis.

#### Protein purification, Western Blot and Co-immunoprecipitation

WT and Ascl1-Venus NSCs were cultured in 10cm diameter dishes, in either proliferation or quiescent conditions for 72h. Media was refreshed after 40h to ensure constant BMP signalling. Cells were then washed with ice-cold PBS, and scraped in Lysis Buffer (ThermoFischer Scientific, 87788) + 1x Protease inhibitor cocktail (ThermoFischer Scientific, 87786) + 1 × EDTA (ThermoFischer Scientific, 87788) + 1x Phosphatase inhibitor cocktail (ThermoFischer Scientific, 78420). Cells were lysed at 4°C for 20min under rotation and then centrifuged at 13,000 RPM at 4°C for 20mins and the pellet discarded. The supernatant was analysed either by western blot or subject to immunoprecipitation.

For western blot analysis, the supernatant was mixed with 1x Laemmli sample buffer (Sigma, S3401-10VL) and incubated at 95°C for 5 mins.

For immunoprecipitation experiments, antibodies were added to cell lysate supernatants and incubated at 4°C for 2 hours under rotation. As controls, mouse anti-V5-tag or rabbit anti-HA-tag antibodies were used under the same conditions. Sepharose coupled to protein G (Sigma, P3296) was blocked with 5% BSA-PBS for 2 hours at 4°C under rotation. After several washes with PBS, it was then added to the lysate-antibody suspension and incubated for 2 hours at 4°C under rotation. After this period, Sepharose beads were washed with lysis buffer 5 times, then suspended in an equal volume of Laemmli sample buffer and incubated at 95°C for 5 mins.

Samples were run in polyacrylamide gel at 120V, after which they were transferred onto a nitrocellulose membrane. Filters were then saturated with 5% BSA in TBS-Tween or 5% milk TBS-Tween and incubated with the antibodies. Detection was performed using ECL Western Blotting Reagents (Sigma, GERPN2106).

### Quantification and Statistical Analysis

To measure immunofluorescence intensity, the nucleus of each identified RGL was manually outlined based on DAPI staining, and the average pixel value of the channel of interest was measured using FIJI software. Every value was normalised to the background level measured in a negative nucleus in the same z-plane as each RGL. At least 200 RGLs in each of at least 3 mice were quantified for each protein. For *in vitro* IHC quantification, average pixel intensity for each channel was measured for the area of each nuclei, using FIJI software. For each experiment, at least 100 cells were quantified across at least 3 biological replicates. To generate the arbitrary units (A.U.) for both *in vivo* and *in vitro* IHC, all the values within a sample were made relative to the average of the control, and multiplied by 100. For quantification of RNAscope^®^ staining, the number of ‘dots’ in each identified RGL nucleus were counted for each probe. In addition, the average pixel intensity in and around each RGL nucleus was measured for each probe, using FIJI. 100 RGLs were quantified across 5 mice. For analysis of Id4 and E47 nucleofected cells, Id4+ or GFP+(E47) cells were identified by immunostaining for Id4 or GFP respectively, and positive cells compared to negative, non-transfected cells within the same coverslip. Cell counts were done from at least 3 coverslips from 3 biological replicates.

For quantification of WB and IP assays, films were scanned and, if appropriate, subjected to band densitometry and quantification using Image J software. Each band value was normalised according to the background of the filter and its loading control.

Statistical analyses were conducted using a two-sample unpaired t test assuming Gaussian distribution using Prism software. All error bars represent the mean ± SEM. Significance is stated as follows: p>0.05 (ns), p<0.05 (*), p<0.01 (**), p<0.001 (***), p<0.0001 (****). Statistical details of each experiment can be found in the figure legend. n represents number of independent biological repeats.

## Data and Software Availability

The accession number for the RNA sequencing data reported in this paper is GEO: GSE116997. If you would like access to this data, please contact the corresponding authors.

## AUTHOR CONTRIBUTIONS

The study was conceived and the manuscript was written by I.B., F.G and N.U. Most of the work was performed by I.B.; B.R. and E.H. provided the Id4 cKO mice and were involved in discussions of the results, M.d.M.M. performed the immunoprecipitations and western blot analyses and E.M., S.V. and D.v.d.B. analysed the RNA-Seq data.

## ACKNOWLEDGMENTS

We gratefully acknowledge Lan Chen for technical support, Rekha Subramaniams and Nicholas Chisholm for managing the mouse colony, the Advanced Sequencing Facility, Bioinformatics and Biostatistics Facility, Experimental Histopathology Facility and Flow Cytometry Facility of the Francis Crick Institute for their help and support, Jane Johnson, Ryoichiro Kageyama, Ana Lasorella, Randall Reed, Jane Visvader for providing mice, Gilbert Rahme, Matthew Havrda and Mark Israel for plasmids, Runrui Zhang and Verdon Taylor for sharing data and manuscript before publication and members of the Guillemot lab for discussions. N.U. was supported by a fellowship from the Francis Crick Institute and is currently supported by the Austrian Academy of Science; B.R. was supported by H2020-MSCA-IF-2014; D.v.d.B. was supported by a Marie Curie Fellowship, project 799214; E.H. was supported by Ligue Nationale contre le Cancer, Fondation ARC pour la recherche sur le cancer (PJA 20131200481, PJA 20151203259) and FP7 Marie Curie CIG. This work was supported by the Francis Crick Institute, which receives its funding from Cancer Research UK (FC0010089), the UK Medical Research Council (FC0010089) and the Wellcome Trust (FC0010089), by the UK Medical Research Council (project grant U117570528 to F.G.) and by the Wellcome Trust (Investigator Award 106187/Z/14/Z to F.G.). The authors declare no conflict of interest. Supplement contains additional data.

